# Unique structural and ligand binding properties of the *Staphylococcus aureus* serine hydrolase FphE

**DOI:** 10.1101/2025.10.12.681921

**Authors:** Jeyun Jo, Tulsi Upadhyay, Xiangyan You, John M. Bennett, Hyunbin Lee, Matthew Bogyo, Matthias Fellner

**Author notes:** Corresponding authors. (M.B.); (M.F.). These authors contributed equally to this work (Jeyun Jo, Matthias Fellner).

## Abstract

*Staphylococcus aureus* is a human pathogen capable of forming biofilms that complicate treatment and facilitate chronic infections. A family of *S. aureus* serine hydrolases are important regulators of virulence and biofilm formation. Among these, FphE is highly specific to *S. aureus* and therefor a viable target for both imaging and therapy. Here, we present bioinformatic and structural evidence that FphE may be involved in aromatic compound metabolism. In addition, twelve distinct crystal forms reveal that FphE exists as a highly unusual but stable and flexible, cross-subunit homodimer, unique to the large alpha/beta hydrolase superfamily. Substrate engagement favors retention of the dimeric state, which is a more catalytically active form of the enzyme and small angle X-ray scattering (SAXS) confirms the dimeric architecture occurs in solution. High-resolution co-crystal structures of FphE covalently bound to two chemically distinct ligands reveal different modes of active site engagement, supporting an atypical structural plasticity of the dimer interface. Together, these findings establish FphE as a structurally unique alpha/beta hydrolase and provide a foundation for structure-guided development of *S. aureus* specific inhibitors and imaging probes.

## Introduction

Worldwide, *Staphylococcus aureus* (*S. aureus*) causes a variety of diseases ranging from local skin or soft tissue infections to invasive chronic infections such as bacteraemia, pneumonia, or endocarditis (*1, 2*). Chronic infections are often linked to the ability of *S. aureus* to form layers of cells embedded within a biomolecular matrix, known as a biofilm (*3, 4*). These robust bacterial structures can form on synthetic surfaces (prosthetic implants) or heart valves, making infections more complicated and costly to monitor and treat (*5*). Biofilms also offer *S. aureus* a relatively protected environment from antimicrobial agents and the host immune response (*6*). As *S. aureus* remains a major human health threat, new methods for rapid detection and therapeutic response monitoring are needed.

Activity-based probes (ABPs), which incorporate weak electrophiles to selectively react with activated amino acid residues, enabled global mapping of active enzymes across diverse biological systems *in situ* (*7*). Our recent study applying a broad-spectrum fluorophosphonate (FP)-based ABP in live *S. aureus* facilitated the discovery of a previously uncharacterized family of serine hydrolases, termed fluorophosphonate-binding hydrolases (Fphs) that are involved in biofilm formation and virulence (*8–18*). Serine hydrolases belong to the alpha/beta hydrolase superfamily, with the serine hydrolase subfamily themselves being one of the largest enzyme families distributed across all kingdoms of life (*19*). They are highly druggable and execute a wide range of biological functions via hydrolytic substrate cleavage of ester, thioester, or amide bonds, allowing them to act as proteases, peptidases, lipases, esterases and amidases. Cleavage is achieved via an active site triad made up of Ser-His-Asp/Glu containing the highly nucleophilic serine residue (*20*). This serine can be targeted by activity-based probes containing a reactive electrophile to label the active enzyme *in vitro*, in cells and even in whole organisms (*7*).

FphE (SAV2581 or Hy30) is a 31 kDa serine hydrolase that lacks homologs in humans or other mortality-associated bacteria aside from *Listeria monocytogenes* (*10, 14*). FphE has emerged as a promising target for the development of imaging probes that can specifically label *S. aureus* via covalent binding to the active site serine. We previously identified triazole urea-based covalent inhibitors for several Fphs, among them FphE (*8*). Optimization of an initial hit resulted in a fluorescent activity-based probe that showed FphE-specific labelling that was highest in cells growing under biofilm-forming conditions (*8*). In addition, FphE levels were found to be dynamic and subject to the bacterial growth state in a strain specific manner.

We have also discovered a potential link between FphE and FphH as activity of FphE was substantially increased in an *fph*H transposon insertional mutant (*8, 17*). The biological role of FphH has not been established but links to stress responses and components of the ribosome rescue system have been described (*11*). FphE can be covalently targeted by β-lactam probes (*21*) and by a carmofur-based probe (*22*). Recently, screening of an oxadiazolone-based library and subsequent lead molecule optimization identified a novel antibiotic compound, named “compound 3” that is active against various *S. aureus* strains (*23*). This compound was shown to covalently bind to a set of ten *S. aureus* proteins, which included FphE and FphH, however, antibiotic potency was not linked to the ability of the compound to bind to the Fphs.

We recently described development ABPs for FphE using the oxadiazolone electrophile of compound 3 as a starting scaffold (*14*). These oxadiazolone-based ABPs were highly specific for FphE and validated it as a target for the development of imaging contrast agents for the rapid and selective detection of *S. aureus* infections. These studies confirmed that FphE is intracellularly located and can be targeted in live bacteria cells without causing cytotoxicity. Furthermore, we demonstrated that a fluorescently labeled version of an oxadiazolone probe can be used to non-invasively image *S. aureus* infection sites in mice.

X-ray crystal structures of FphE in the unbound form or with the oxadiazolone covalently bound in the active site demonstrated a highly unusual inter-subunit homodimer where the catalytic triad of each monomer is composed of components of each of the two monomers (*14*). Interestingly, we recently found that FphE inhibitors bearing a boron warhead generate a covalent tetrahedral adduct with the catalytic serine from one monomer and the histidine from the other monomer, thereby preventing dimer dissociation of FphE (*18*).

Here we present detailed structural features and the molecular basis for ligand selectivity of FphE. Our bioinformatic analysis reveals FphE is genomically linked to enzymes predicted to process aromatic substrates, providing a rationale for its preference for aromatic probes. Biochemically, we demonstrated that FphE forms a homodimer that has increased catalytic activity and is stabilized upon substrate engagement. Through the analysis of twelve distinct crystal forms including co-crystal structures with multiple ligands, we demonstrate an overall structural plasticity of this unique homodimer. We also identify two distinct binding pockets including the previously reported hydrophobic tunnel and an adjacent pocket near the catalytic site. While this unprecedented dimer architecture enables ligand-dependent shifts in binding modes, bioinformatic analysis and our co-crystal structures reveal that selectivity is dictated not by the conserved active site, but by engagement of the less-conserved, peripheral hydrophobic tunnel. Together, these insights provide a molecular rationale for FphE selectivity and establish a framework for the rational design of *S. aureus*-specific probes.

## Results

### Taxonomic distribution and genomic context of FphE suggest association with aromatic compound metabolism

To understand the biological role of FphE, we searched for homologs using a position-specific iterative BLAST (PSI-BLAST) querying FphE sequence against the National Center for Biotechnology Information (NCBI) non-redundant protein database (fig. S1 and Supplementary Data 1). To ensure high structural and sequence similarity while minimizing redundancy, we excluded sequences with <40% identity and clustered the remaining 1,619 proteins at a 95% identity cutoff, yielding 119 representatives as a final set. Interestingly, all proteins are encoded within species from the phylum *Bacillota* with *Listeria monocytogenes* (N = 5) being the only mortality-associated species (*24*), apart from *S. aureus* (N = 12; Supplementary Data 2). This highly restricted distribution highlights the potential of FphE to serve as a selective imaging and diagnostic target for *S. aureus* infections (*14*). The closest homologs outside *Bacillota* are mainly encoded by fungi, suggesting a specialized role for this protein within the phylum (fig. S1). We found that *fphE* is conserved across diverse *S. aureus* strains under distinct gene names (e.g., SAUSA300_2518, strain USA300; NWMN_2480, strain Newman; SA2367, strain N315; SACOL2597, strain COL; SAR2661, strain MRSA252; HMPREF0769_10569, strain MN8; SAV2581, strain Mu50/ATCC 700699) and that its homologs are differentially regulated under various environmental, metabolic, and host-associated stresses (*17, 25–37*), although the underlying functional relevance of this regulation remains unclear (table S1 and Supplementary Data 3). In *L. monocytogenes*, transcription levels of *fphE* were increased in a nonvirulent strain and in a virulence factor knockout (Δ*prfA*), indicating a potential role in pathogenicity (*38–41*). These findings collectively suggest that FphE is a taxonomically restricted enzyme closely related to stress-related pathways in *S. aureus*, supporting its potential as a species-selective biomarker.

To further explore the biological function of FphE, we examined its and its homologs’ genomic loci to identify possible conserved pathway associations (Fig. 1, A and B, and fig. S2). In *S. aureus* USA300, *fphE* (SAUSA300_2518) is encoded in a seven-gene operon (SAUSA300_2512–2518) associated with biofilm regulation (Fig. 1A) (*42*). This operon is divergently transcribed into two mRNAs (SAUSA300_2512–2514 and SAUSA300_2515–2518), with the transcription of both controlled by a TetR-family repressor GbaA (SAUSA300_2515; glucose-induced biofilm accessory protein A) binding to the shared operator (Fig. 1A) (*42*). The covalent modification of GbaA by electrophiles (*N*-ethylmaleimide and methylglyoxal) induces these operons, and a double mutant lacking both *gbaA* (also known as *rob*) and *fphE* showed no additional effect on biofilm formation compared to the *gbaA* single mutant in *S. aureus* strain FK300 (*42, 43*). Comparative genomic analysis revealed that the latter four genes of the operon that include *fphE* (SAUSA300_2515–2518) are almost exclusively conserved across *Staphylococcus* species (23 out of 55 representative loci) with only one example outside the genus *Staphylococcus* (in *Macrococcus equipercicus* ATCC 51831; Fig. 1B and fig. S2). Interestingly, we found that *fphE* and an upstream gene for an amidohydrolase (SAUSA300_2517) are genomically co-localized in a majority of the representative loci across *Staphylococcus* and other bacterial species, suggesting possible cooperation within a biochemical pathway (80 amidohydrolases in 59 out of 119 analyzed loci; Fig. 1B and fig. S2). Moreover, in two *S. aureus* strains (isolate P15 and st1226), *fphE* and the upstream amidohydrolase gene are fused as a multidomain protein (RefSeq accessions HCY5833117.1 and CXW27440.1; fig. S2).

**Fig. 1.**
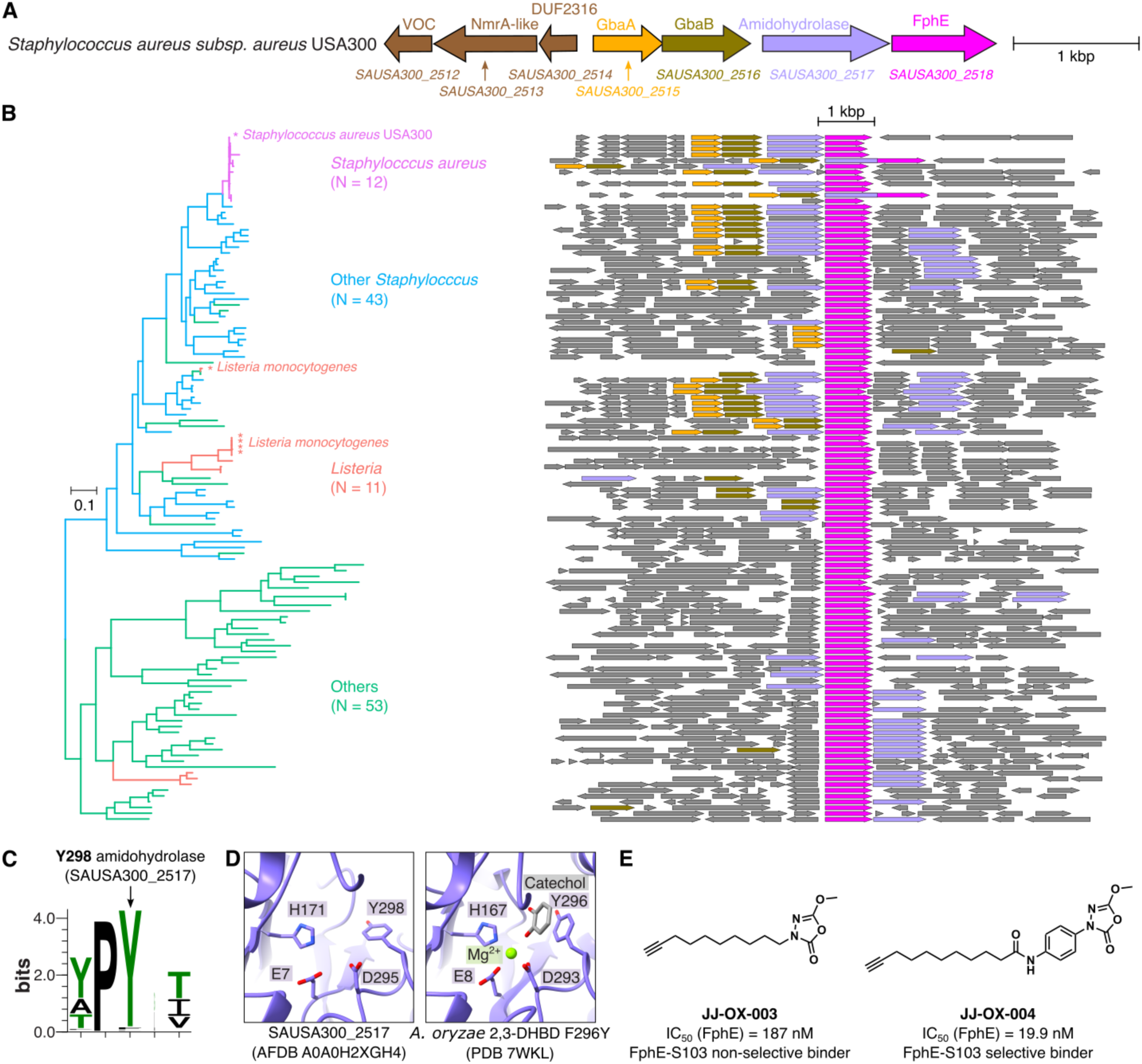
FphE is associated in aromatic compound metabolism. (**A**) Genomic locus of the *S. aureus* USA300 *fphE* (SAUSA300_2518). Protein names or predicted domains are given above and gene names are given below arrows. (**B**) A phylogenetic tree of 119 representative FphE homologs. Genomic loci are shown, and arrows are colored by their homology. Names of all species are shown in fig. S2. (**C**) A sequence logo for Y298 of the amidohydrolases (SAUSA300_2517) and 79 homologs encoded near genes for FphE homologs in (B). (**D**) Active sites of the AlphaFold2-predicted amidohydrolase (SAUSA300_2517) (left) and *A. oryzae* 2,3-dihydroxybenzoate decarboxylase (2,3-DHBD) F296Y (PDB 7WKL; right) (*44, 76*). (**E**) Inhibition potency and selectivity of JJ-OX-003 and JJ-OX-004 toward the catalytic serine (Ser103) of FphE (*14*).

Sequence alignment of the aminohydrolase SAUSA300_2517 and its 79 homologs identified across 50 of the 119 loci revealed a highly conserved tyrosine residue (Y298), along with putative metal-binding residues (E7, H171, and D295; Fig. 1C and fig. S3A). Structural homologs (PDB: 7WKL, 7WMB, 6JQX, and 7BPC) of the AlphaFold model of SAUSA300_2517 utilize corresponding aromatic residues (tyrosine or phenylalanine) to engage π-π interactions for recognizing the benzene ring of their aromatic substrates, such as catechol, salicylic acid, and 2,5-dihydroxybenzoic acid (Fig. 1D, fig. S3 B–D, and Supplementary Data 4) (*44–46*). Consistent with this, in our previous work, we observed that FphE was efficiently labeled by probes that contain an aromatic ring (*14*). In particular, the probe JJ-OX-004 that contains an aromatic ring between the lipid chain and oxadiazolone warhead showed selectivity toward the catalytic serine (Ser103) of FphE with dramatically improved potency compared to JJ-OX-003, which lacks this aromatic feature (Fig. 1E). We previously demonstrated that FphE has carboxyesterase activity against fluorogenic substrates, with preference for 4-methylumbelliferyl caprylate (4-MUC) (*14*). In the present study, we further show that FphE lacks protease activity (table S2). Taken together, these observations suggest that FphE as an esterase and the upstream amidohydrolase may participate in a shared biochemical pathway for modifying aromatic substrates.

### FphE forms a homodimer in solution

FphE has a predicted molecular weight of 31 kDa and SDS-PAGE analysis of probe labeled *S. aureus* cell lysates, as well as recombinantly produced FphE, confirm that the protein resolves at this predicted molecular weight (*8, 14, 17*). However, when recombinant FphE was subjected to size exclusion column chromatography, two distinct but relatively equal peaks were observed that resolve as the same molecular weight (∼31 kDa) when analyzed by SDS-PAGE analysis (Fig. 2A). We subjected the purified protein peak 1 and peak 2 to native-PAGE under nonreducing conditions and observed two distinct protein bands, a dimer corresponding to peak 1 and a monomer corresponding to peak 2 (Fig. 2B). We observed no interconversion between the monomer and dimer upon changing the concentration of either form. Although each form contained a small amount of the other as a contaminant, no change in equilibrium was observed when the samples were progressively diluted to lower protein concentrations and directly analyzed by native-PAGE. We tested whether labeling by our FphE-selective probe JJ-OX-007 containing a BDP-TMR fluorophore (*14*) affects the oligomeric state of FphE (Fig. 2C), as well as examined whether the oligomeric status could be modulated by addition of the lipid ester substrate, 4-MUC. We incubated the dimeric and monomeric form of FphE at room temperature for 30 minutes in the presence or absence of substrate, and the samples were either treated with JJ-OX-007 or left untreated and incubated for an additional hour at 37 °C (Fig. 2D). Interestingly, we observed spontaneous conversion to the monomeric form (Fig. 2E), that was reduced by the co-incubation with the 4-MUC substrate. (Fig. 2E, lanes 1 and 2, Coomassie stain). A similar trend was observed upon additional treatment with the JJ-OX-007 fluorescent probe that enabled more sensitive detection of both monomeric and dimeric forms compared to Coomassie staining (Fig. 2, E and F, and table S3). We also evaluated the effects of temperature on dimer-to-monomer conversion (fig. S4). To minimize effects during substrate incubation time, we transferred the samples to the target temperatures after substrate and probe addition. As expected, the presence of the substrate strongly prevented the conversion of dimer to monomer at 37 °C (fig. S4, top row). However, at lower temperatures such as room temperature or 4 °C, no change in oligomeric status was observed even after 1 hour incubation. Fluorescence from probe labeling decreased for both forms as the temperature decreased, which is likely due to overall reduced enzyme activity at lower temperatures (fig. S4, left column). Protein from both size exclusion peaks generated the same crystal forms. However, the dimer showed higher cleavage activities towards 4-MUC model substrate than the monomer (fig. S5).

**Fig. 2.**
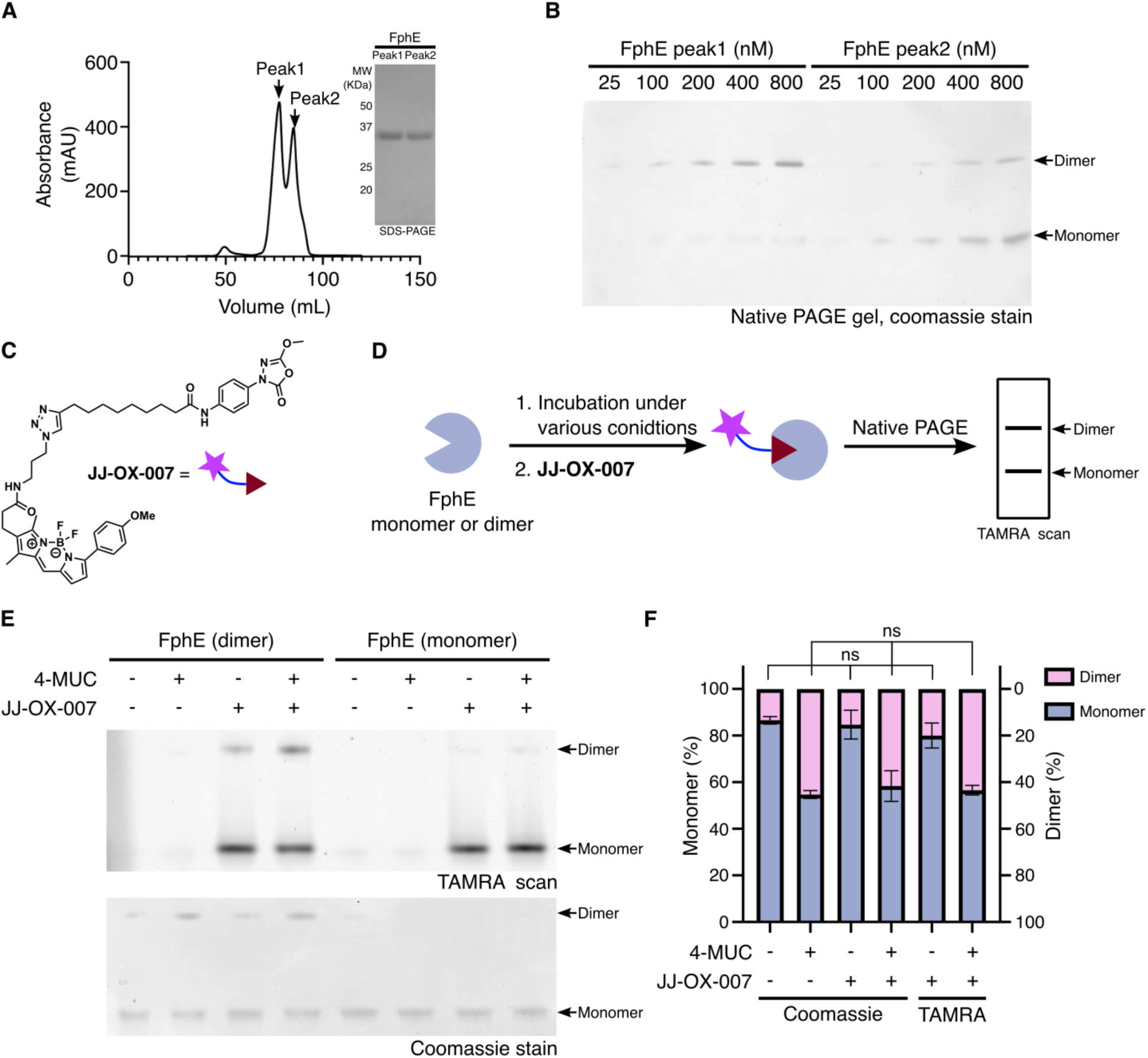
Biochemical characterization of FphE oligomeric states and probe labeling. (**A**) Size exclusion chromatography of recombinant FphE revealing two major peaks corresponding to dimeric (peak 1) and monomeric (peak 2) forms. (**B**) Native-PAGE analysis of increasing protein concentration from each peak. (**C**) Chemical structure of FphE covalent fluorescent probe JJ-OX-007. (**D**) Schematic overview of fluorescent probe-based detection of monomeric and dimeric FphE. (**E**) Native-PAGE analysis of oligomeric status and probe labeling of FphE monomer and dimer. (**F**) Quantification of the native-PAGE in Figure 2E. Values are shown as mean ± SD (*n* = 2) from independent experiments. 4-MUC, 4-methylumbelliferyl caprylate; ns, not significant.

### FphE crystals assemble into an unusual homodimer architecture in 12 unique crystal forms

FphE crystals grew in various conditions into a range of different crystal forms (table S4). FphE crystallized in a pH range from 5.5 to 8.5 in the presence of 15-30% polyethylene glycol compounds or in high salt conditions (table S4, S5 and S6). Crystal growth was accelerated by the presence of magnesium, potassium and calcium salts but crystals also appeared in the absence of any additional salts. Processing revealed 12 distinct crystal forms of FphE. We arranged them roughly by their highest diffracting crystal. Crystal form 1 and 12 were the most frequently appearing crystals with form 1 always diffracting to high resolution (as high as 1.38 Å) and form 12 crystals never diffracting to better than 2.5 Å and often worse than ∼5 Å. The form 12 crystal could not be phased using molecular replacement due to severe tNCS (translational noncrystallographic symmetry). Interestingly, these 1 and 12 crystal forms sometimes appeared in the same crystallization drop and they could not be distinguished based on optical presentation. The remaining 10 crystal forms appeared far less frequently and were sometimes also found in the same crystal drop as form 1 and 12. Occasionally, their occurrence appeared to be correlated with the presence of specific compounds in the reservoir mixture (table S4) or when specific ligands were co-crystallized, but no concrete pattern could be established. Crystal form 10 could also not be processed due to potential twinning issues causing complications at molecular replacement. Crystal form 11 suffered from poor diffraction patterns that prevented processing beyond merging and scaling steps. The remaining 9 crystal forms of FphE were fully processed, refined and deposited to the Protein Data Bank (*47*) (table S7 and S8).

Although hundreds of FphE crystals were processed, the monomeric form of FphE was never observed in the crystalline state with all crystal forms being a homodimer (Fig. 3). In all cases the homodimer is formed in a very unusual way where the N-terminal residues 1–136 of one FphE monomer are assembled with the C-terminal residues of 168–276 of the second FphE monomer to create two, connected “full-length” FphE molecules that may resemble the monomeric state. These monomers are connected via the remaining residues 137–167 which form a long bent helix that runs antiparallel to the corresponding helix from the other FphE monomer (Fig. 3A). The active site still contains the conserved catalytical triad for serine hydrolases, in this case formed by Ser103, His257, and Glu127, with Ser and Glu originating from one FphE monomer and His from the other. The hydrophobic active site pocket and access tunnel (Fig. 3B–D) are therefore also formed by residues from both FphE monomers. Specifically, the connecting helix ∼155–167, as well as helices 170–178 and 183–207 contain residues from both monomers. These helices form a so called “lid” in hydrolases of a similar fold (*48*), including FphH (*11*) or FphI (*15*). However, the FphE conformation is rather unique in that the helices form a distinct entrance to the active site rather than a dynamic lid that may cover the active site after ligand binding. Our previous prediction of the monomer of FphE (*9*), as well as predictions using Alphafold2 (*49*) for this study, suggest that FphE has two helices in place of the long connecting dimer helices 136–154 and 156– 167 (Fig. 3D). It appears that the dimer is formed when the C-terminus starting at residue 155 bends away from the core protein with residues 156–167 forming a long helix instead of a helix that bends backwards towards the FphE monomer. However, the bend at the middle of the long connecting dimer-helix is still retained to some degree in all crystal forms and appears to be the main differentiating factor of the crystal forms. Cα alignment of the first ∼1–136 residues of one FphE monomer demonstrates how the connecting helix differs significantly between the crystal forms (Fig. 4 and fig. S6). Alignment of the N-terminus results in the ends of the connecting helices (distance of Cα atom of residue 137 and 167) being between 13.1 to 19.2 Å across the different crystal forms. This is due to the extent of bending in the connecting helices which is characterized by a shift of the central residue 155 of the helices. Due to the differences in the bend of this linker region (table S9), the positions of the two assembled FphE monomers shift relative to each other. Alignment of one FphE copy, shifts the other one by up to 25 Å in one axis and by up to 13 Å in another axis across the different crystal forms. The shifted positions of the two FphE monomers then facilitate variable crystal interfaces which results in the multiple crystal forms. These unique folds for the various helices can be influenced by ligands covalently bound to the active site serine. Some crystal forms were only observed in the presence of a specific ligand: form 2 only with oxadiazolone compound 3, form 4 and 6 only with borolane-based compound Q41 and form 7 only with boronic acid-based compound W41 (*18*). The different crystal forms also influence the accessibility of the active site (fig. S7), with all crystal forms showing a similar open hydrophobic tunnel leading to the active site but with slight variations in the dimensions and arrangement of the tunnel.

**Fig. 3.**
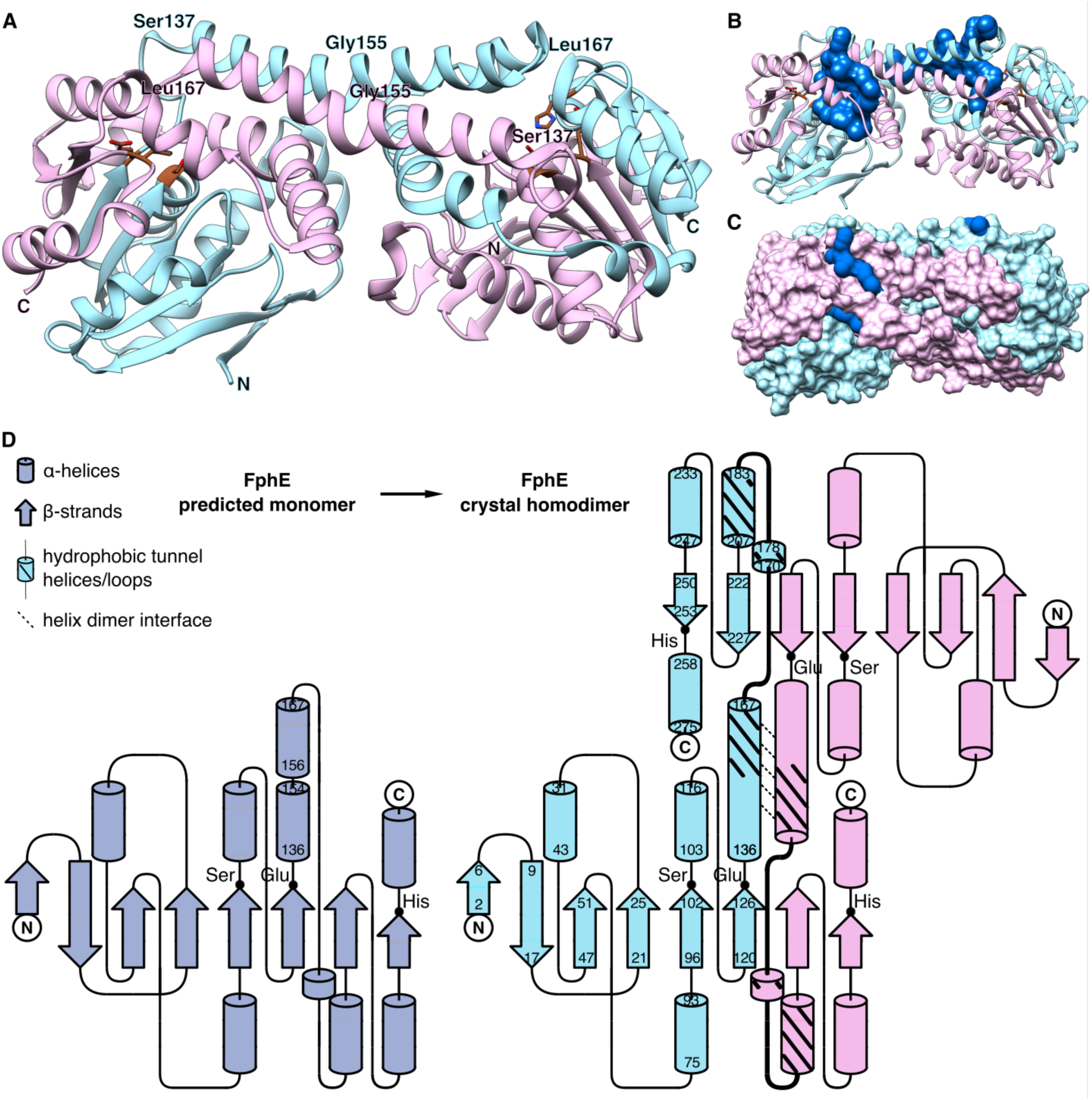
FphE homodimer. (**A**) Ribbon representation of FphE homodimer in crystal form 1 (PDB ID 8T87, 1.62 Å) (*14*), FphE chain A in light cyan and chain B in pink, with active site triad Ser-His-Glu displayed in brown. N and C termini annotated; dimer interfacing helix residue numbers indicated. (**B**) Active site pockets illustrated as a blue surface model, identified via the FPocketWeb browser (*98*). (**C**) Protein chains and active site pockets surface representation. (**D**) Secondary structure schematic of predicted monomer in lavender compared to FphE homodimer in crystal form 1 (8T87) with FphE chains in blue and pink. N and C termini annotated with capital letters. Active site triad residue location noted with a black dot. For the homodimer chain A residue numbers for each helix and strand are indicated. Hydrophobic tunnel helices and loops marked with thick black lines. Helix dimer interface illustrated via dotted lines.

**Fig. 4.**
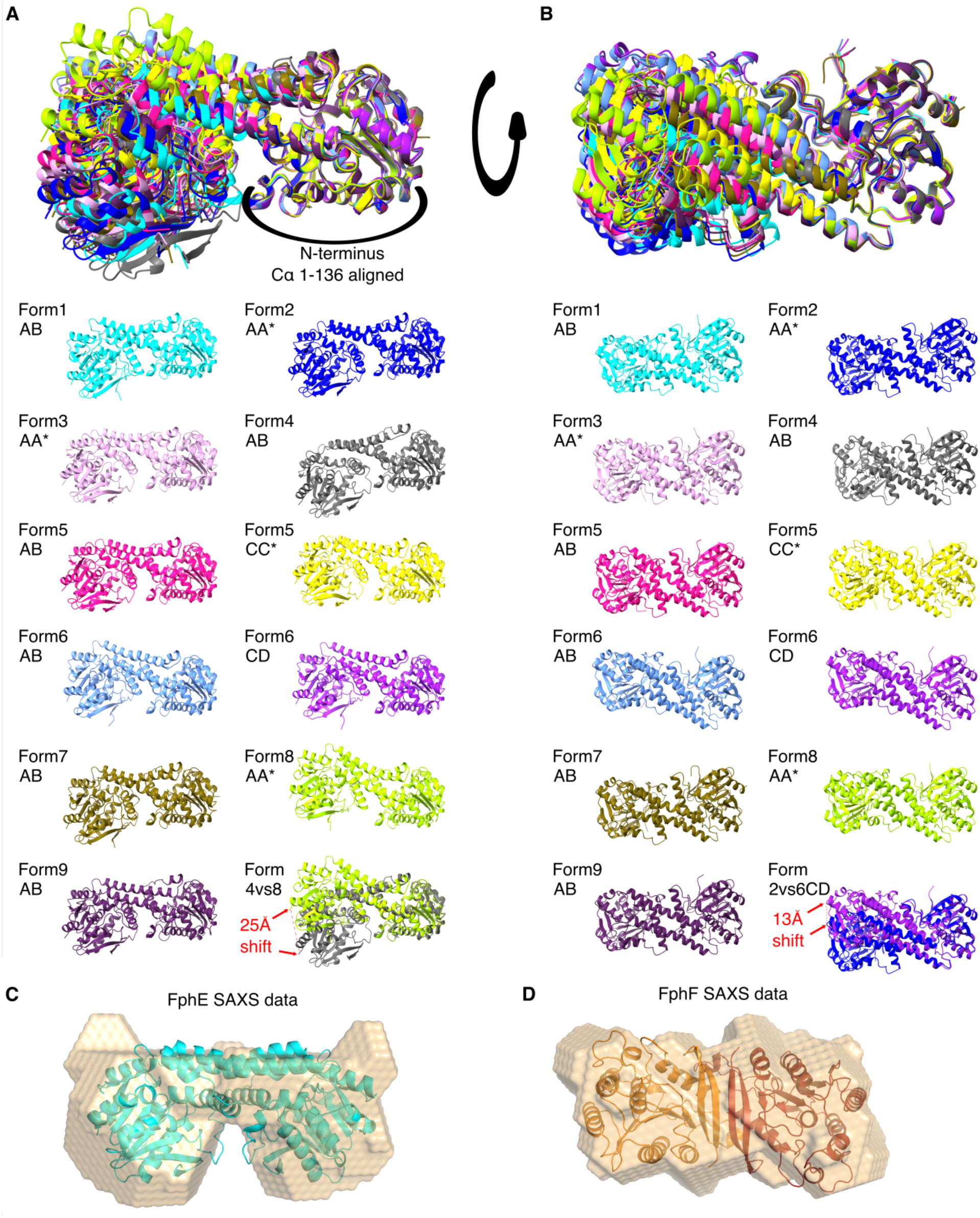
FphE crystal forms. (**A**) Ribbon representation of alignment of chain A (or C in two cases) Cα atoms of N-terminal residues 1-136 of each crystal form. Overlay of all crystal forms and individual pictures for each crystal form in different ribbon colors. Form 1 chain A+B dimer cyan (PDB ID 8T87, dataset resolution 1.62 Å) (*14*); form 2 A+A* crystal symmetry dimer blue (8G49, 1.60 Å); form 3 AA* pink (8G48, 1.95 Å); form 4 AB grey (8UWM, 1.97 Å) (*18*); form 5 AB magenta, CC* yellow (9COM, 2.04 Å); form 6 AB light blue, CD purple (9EBF, 2.50 Å); form 7 AB ochre (8UIX, 2.39 Å) (*22*); form 8 AA* green (9D87, 2.30 Å) and form 9 AB dark purple (9EDJ, 2.60 Å). Largest shift of other dimer end between crystal form 4 and 8 of 25 Å (C-terminus residue 276 Cα atom) indicated in red. (**B**) Same alignment turned by ∼90 degrees. Largest shift of other dimer end between crystal form 2 and 6 of 13 Å (residue 247 Cα atom) indicated. (**C**). Small-angle X-ray scattering (SAXS) result of FphE homodimer. Ab initio envelope (beige surface model) of FphE in solution overlayed via ATSAS-CIFSUP with FphE crystal form 1 (8T87). (**D**). SAXS result of FphF homodimer. Ab initio envelope of FphF in solution overlayed via ATSAS-CIFSUP with FphF crystal (6VH9, chain A in red and chain D in orange).

In some cases, the two monomers of FphE that form the homodimer are identical, resulting in a crystal form with only one chain A with an identical crystal symmetry mate A* forming the homodimer (table S4, form 2, 3 and 8). In other cases, the homodimer is formed via chain A and chain B of the asymmetric unit (form 1, 4, 7 and 9). In one case the asymmetric unit contains two homodimers of chain A with chain B and a second one formed by chain C and chain D (form 6). And the final form contains a homodimer formed by chain A with chain B with a second homodimer formed by chain C and its crystal symmetry mate C* (form 5). It was especially challenging to obtain the correct molecular replacement solution for this crystal form without prior knowledge of the correct assembly.

The long antiparallel connecting helices of residues 136–167 set up a domain with hydrophobic residues Ile147, Val148, Ile151, Leu152, Leu156, Met160, Phe163 residues involved in the dimer interface and charged residues Thr138, Lys141, Asp142, Asp145, Asp146, His149, Thr153, Glu154, Glu157 and Lys161 facing the opposite, solvent exposed, direction. This hydrophobic interfacing and hydrophilic solvent exposed arrangement is a potential key driver of the unusual helix conformation of the dimer. Our sequence conservation analysis also indicates that the hydrophobic interfacing residues are highly conserved in the 119 FphE homolog sequences, with the non-interfacing residues showing less conservation, while mostly retaining their charged state (fig. S8 and Supplementary Data 5). Ile151 is the least conserved among the hydrophobic interface residues, but it is frequently replaced with either valine or glycine (Supplementary Data 5). It is plausible that other alpha/beta hydrolase enzymes with a similar conservation pattern at this region of the fold may also be able to adopt the FphE dimer configuration.

### The FphE homodimer crystal architecture is present in solution

While we were able to confirm that FphE crystalizes as a dimer and also resolves in solution as a dimer by size exclusion chromatography, we wanted to determine how the structure of the dimer in solution relates to the observed unique crystal architectures. Therefore, we determined the in-solution structure of FphE using Small-angle X-ray scattering (SAXS, table S10). FphE SAXS data (Figs. 4C, S9A, and S10) showed that the dimer peak clearly contained a dimer structure of FphE in-solution that closely matched the crystal structures of FphE. Crystal form 1 was a good match to the SAXS data with the connecting helix clearly visible. Minor deviations of the data from crystal form 1 (fig. S9A) further confirm the flexibility of the FphE homodimer suggested by the multiple crystal forms. Creation of an updated SAXS data-based model of crystal form 1 (fig. S9B) emphasized this flexibility as the overall fold remains a close match (fig. S9C) with only the middle of the connecting helices altered to better match the SAXS data. In addition, we determined the SAXS in-solution structure of FphF. Both are serine hydrolases of similar size, with both adopting a traditional alpha/beta hydrolase fold, yet they do not appear to be functionally related with only 23% sequence identity. We previously described FphF as a homodimer in solution, however the crystal structure showed a tetramer (*9*). We proposed that the FphF homodimer is formed via antiparallel association of its first β-strand. Fitting such a homodimer was the best fit for the SAXS data, confirming our hypothesis. (Figs. 4D, S11A and S12). Creation of an updated model of the FphF crystal structure to fit the SAXS data did not result in significant changes, suggesting a less flexible and more uniform structure of FphF compared to FphE (fig. S11, B and C). While the N-terminal homodimer interface of FphF is uncommon for alpha/beta hydrolases, the FphE interface appears to be completely unique for this enzyme.

### The FphE homodimer is unique within the large alpha/beta hydrolase superfamily

FphE has a general alpha/beta hydrolase fold (Inter Pro IPR029058) (*50*), making it related to a group of over 3,000 proteins containing this fold in the protein data bank (PDB) (*47*). This overall set of structures is made up of subfamilies AB_hydrolase_1 (934 experimental determined structures), AB hydrolase superfamily (191 structures) and Pfam Abhydrolase_6 (PF12697, 179 structures). These groups are very broad and thus do not provide insights into a putative biological function of FphE. We used the FphE Alphafold 2 (*49*) predicted monomer structure (table S11) and dimer crystal structure (table S12 and S13) to screen for 3D matches in the PDB (*47*) as well as in the predicted proteome of *S. aureus* using the DALI webserver (*51*). For the monomer, the top hit was Zearalenone lactonase (*52*) which structurally aligns well in the β-sheet region and some helices (129 Cα pairs at 1.2 Å rmsd) but the active site loops and helices, including lid helices differ significantly and the proteins have only 18% sequence identity. In fact, all of the top hits for the predicted FphE monomer structure have similarly low sequence identity and align structurally in a similar way with decreasing statistics of the overall alignments. Searching for structure matches to FphE in the dimer conformation only resulted in hits that loosely match the first ∼140 residues with statistics indicating that these are only general alpha/beta fold hits. Thus, the unusual FphE dimer conformation appears to lack any matches to proteins in the PDB (*47*) as well as to predicted structures of the *S. aureus* proteome found in AlphaFoldDB. In addition, AlphaFold 3 (*53*) was also unable to predict the FphE dimer that we observed in our determined crystal structures.

### The structure of FphE with a covalent fluorophosphonate-probe reveals shared binding features with JJ-OX-004 despite distinct electrophile chemotypes

FphE is covalently inhibited by electrophilc chemotypes including fluorophosphonates (FPs) (*17*), triazole ureas (*8*), oxadiazolones (*14, 23*), β-lactams (*21*), carmofur (*22*), and boronic acids (*18*). Members of the Fph protein family have been systemically profiled using the activity-based probe FP-tetramethylrhodamine (FP-TMR). To enable further structural analysis of FphE, we designed and synthesized an alkyne version of the FP probe, JB101 (Scheme S1) and then determined the structure of FphE bound to this molecule. As expected, JB101 forms an irreversible covalent bond to the active site Ser103 residue from one FphE dimer copy (residues 1-136), with the fluorine atom of JB101 being replaced with the oxygen side chain of the serine residue (Fig. 5, A to C). The core aliphatic backbone of the ligand is located in the hydrophobic active site tunnel that we described previously in detail (Fig. 5, D and E) (*14*). The phosphoryl oxygen atom of JB101 occupies the well described oxyanion hole (*54, 55*) of alpha/beta hydrolases, formed here via interactions with main chain amide groups of Ser104 and nearby Ala28 from the same chain as the active site serine (Fig. 5, B and F). In the case of FphE, this atom is further stabilized via hydrogen bonds to two side chain orientations of Ser104. There are many additional hydrophobic interactions along the JB101 ligand backbone. Carbon atoms near the phosphor atoms are stabilized by backbone carbons of Gly27 and the Cβ atom of Asn29. All other interactions are formed with the other dimer copy (residues 168-276) of FphE which forms the active site tunnel via the end of the dimer connecting helix 156–166 and the following 170–178 and 183–207 helices, as well as the loop containing the active site triad residue His257 (∼256–261). The electron density quality of JB101 atoms of the alkyl linker decreases with distant to the covalent link to the active site serine, suggesting that the ligand may have alternate conformations in the tunnel. This is a similar ligand binding orientation to what we previously reported for the JJ-OX-004 ligand which contains an oxadiazolone electrophile (fig. S13, A and B) (*14*). In addition, based on our structure, the terminal alkyne group of JB101 is likely able to reach the protein surface, which explains how this probe can easily be modified via click chemistry without losing specificity for FphE.

**Fig. 5.**
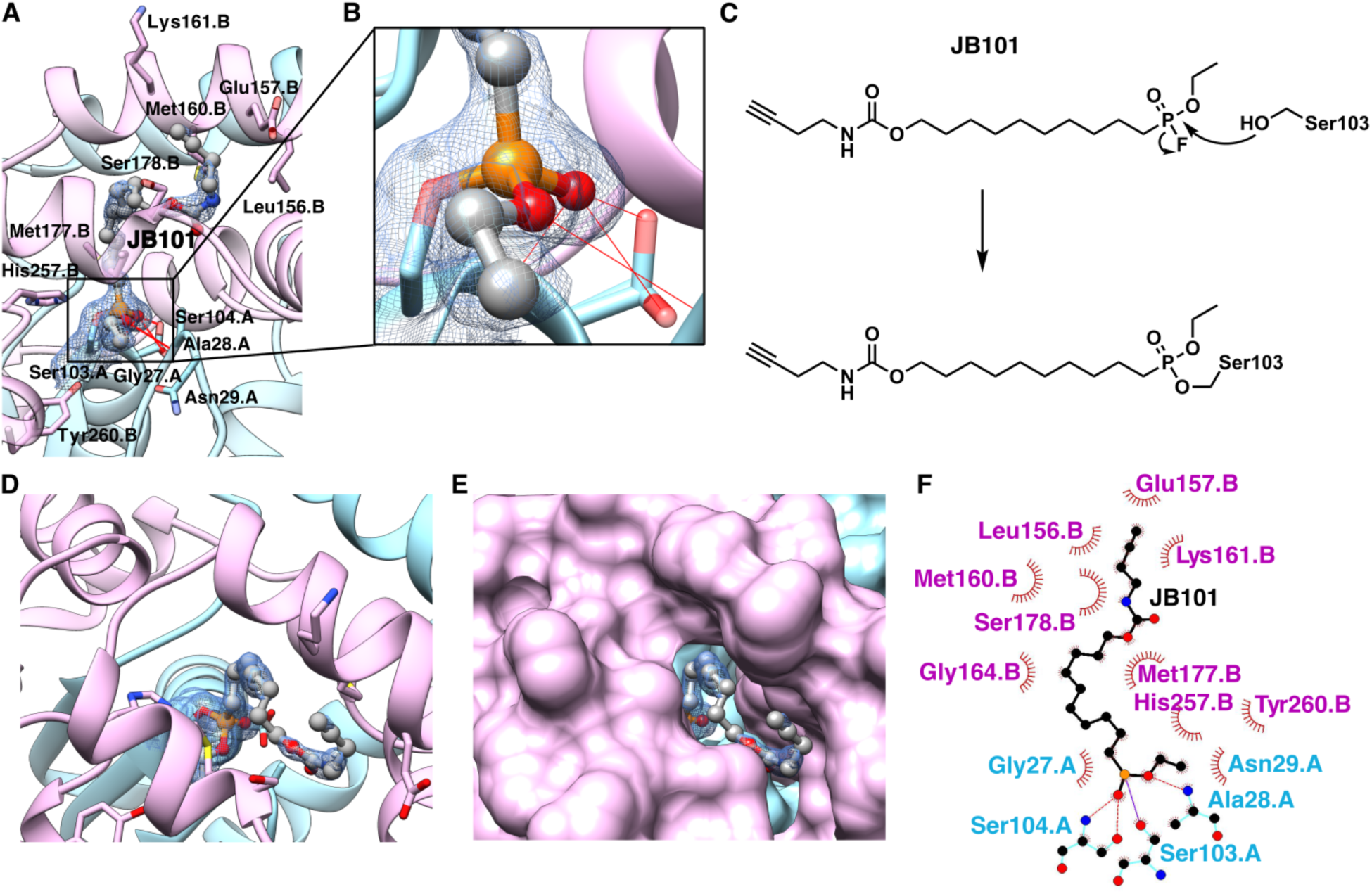
FphE covalently bound by FP-JB101. (**A**) Crystal structure of FphE in complex with fluorophosphonate compound JB101 (crystal form 1, PDB ID 8SBQ, at 1.50 Å). Ribbon representation with carbon atoms of FphE chain A in light cyan, chain B in pink (helix 171.B-178.B in the foreground transparent), JB101 carbon atoms in grey, hydrogen bonds in red. Side chains of key residues displayed and labelled. The 2Fo−Fc electron density map for JB101 and Ser103 of chain A is shown as a blue mesh at 1 σ. (**B**) Closeup of indicated window in (A). (**C**) JB101 chemical structure with nucleophilic attack of active site Ser103 indicated. (**D**) View from above in either ribbon or (**E**) surface representation. (**F**) LigPlot 2D illustration of JB101 interactions with FphE residues.

### The oxadiazolone compound 3 occupies a unique pocket compared to other ligands

We next determined the crystal structure of FphE with an oxadiazolone based molecule, compound 3, that was recently discovered to have antibiotic potency against methicillin resistant *Staphylococcus aureus* (MRSA) (*23*). This inhibitor covalently binds to ten proteins in MRSA bacteria, including FphC, FphE and FphH. Further studies suggested that combined inhibition of FabH (an essential β-Ketoacyl-Acyl Carrier Protein Synthase), FphC, FphE, and AdhE (an aldehyde−alcohol dehydrogenase) contributes to the antimicrobial activity of compound 3. We previously described the inhibition of FphH by compound 3 (*11*) but no structure of this molecule has been reported. Co-crystallization of FphE with compound 3 resulted in crystal form 2 that was only observed in the presence of this molecule. In this form, two homodimer copies of FphE are perfect crystal symmetry mates A and A* (Fig. 6, A and B). As expected, compound 3 is covalently bound to the active site Ser103 with the oxadiazolone ring opened to form a stable carbamate linkage as was proposed previously (Fig. 6C) (*23*). Consistent with the crystal structure of FphE bound to the related oxadiazolone probe, JJ-OX-004, (*14*) the free oxygen atom of the oxadiazolone group is located near the oxyanion hole (*54, 55*) but not directly in the hole (Fig. 6A). The oxygen still forms a hydrogen bond with the amide group of Ala28 but in contrast to JB101 (2.9 Å O-N distance), the atom is shifted to no longer be able to form a hydrogen bond with the Ser104 backbone nitrogen atom (4.2 Å O-N distance). Beyond the oxadiazolone group, compound 3 does not fully occupy the hydrophobic tunnel, and does not reach the surface of FphE as described for JB101 or JJ-OX-004 (fig. S13, A to C). In contrast, compound 3 occupies a nearby pocket formed by residues Ala28, Asn29 located in a loop close to the active site serine, and Arg193, Thr194 and Trp197 from the other dimer copy, located on helix 183–207 (Fig. 6, A and D). In addition, a hydrogen bonding network, with several water molecules, links the ligand to residues Ser102, Asn144 from one FphE dimer copy and Thr194, His257, Tyr260 from the other copy (Fig. 6B). This pocket is covered by helix 156–166 and the following 170–178 helix that connect the monomers in the dimer (Fig. 6E). Hydrophobic interactions are observed with Met160, Phe163, Leu167, Ile169, Met177 and Trp197 (Fig. 6F).

**Fig. 6.**
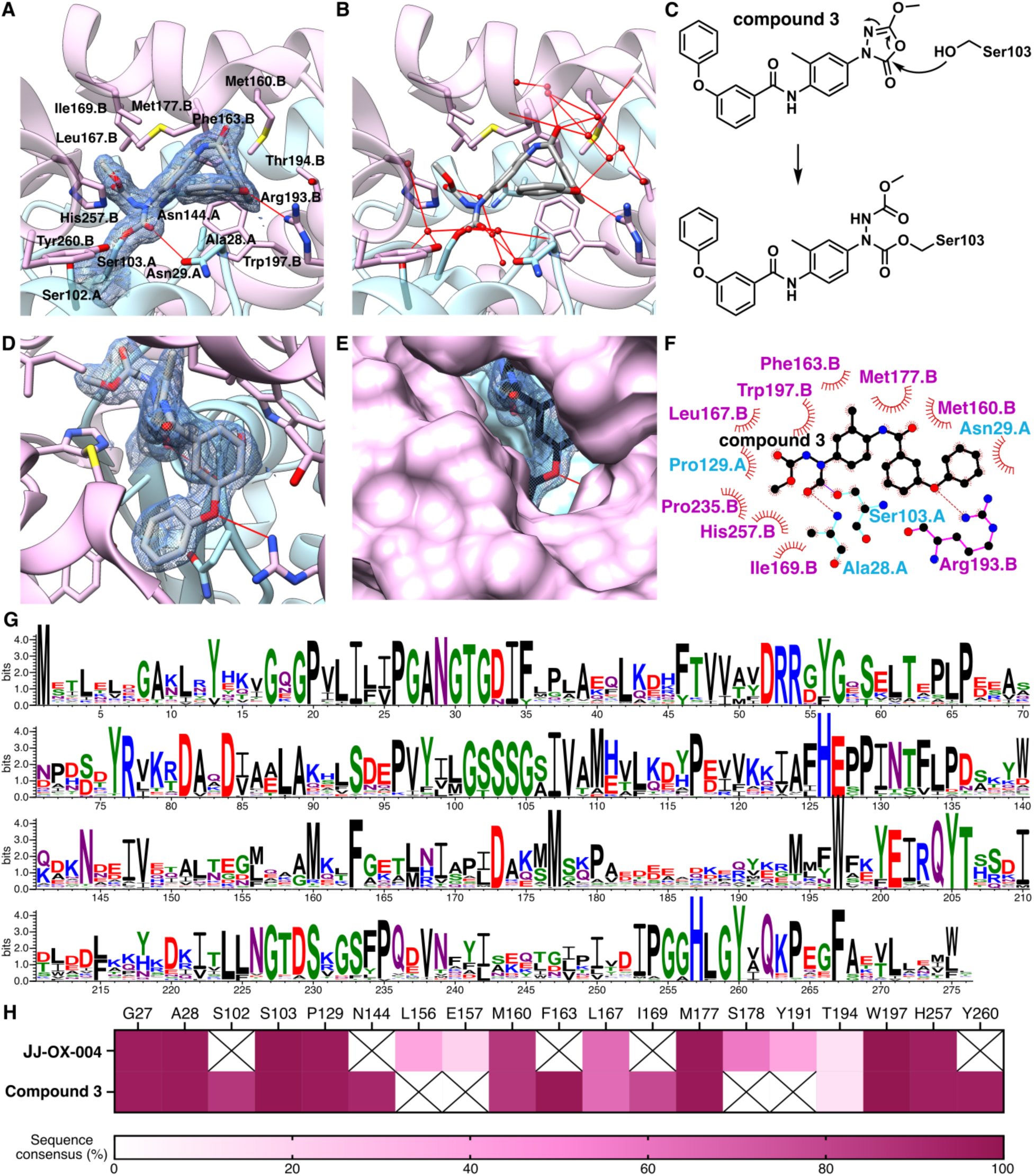
FphE covalently bound by compound 3. (**A**) Crystal structure of FphE in complex with oxadiazolone compound 3 (crystal form 2, PDB ID 8G49, at 1.60 Å). Ribbon (transparent) representation with carbon atoms of FphE chain A in light cyan, chain B in pink, compound 3 carbon atoms in grey. Side chains of key residues displayed and labelled. The 2Fo−Fc electron density map for compound 3 and Ser103 of chain A is shown as a blue mesh at 1 σ. (**B**) Hydrogen bonding network of compound 3 in the same orientation as in (A) depicted with water molecules as red spheres and hydrogen bonds as red lines. (**C**) Compound 3 chemical structure and reaction with active site Ser103 indicated. (**D**) Further rotated view of A, view from above in either ribbon or (**E**) surface representation. (**F**) LigPlot 2D illustration of compound 3 interactions with FphE residues. (**G**) Sequence WebLogo representation of 119 FphE homologs. (**H**) Heatmap of sequence consensus (%) for key residues forming the binding pocket or interacting with JJ-OX-004 and compound 3. Boxes with a cross indicate residues without direct interactions.

### Two distinct binding sites in FphE underlie pocket-specific ligand selectivity

To assess whether ligand binding correlates with evolutionary conservation, we generated sequence logos for 119 FphE homologs using WebLogo (*56*) and calculated residue-wise consensus values for positions interacting with both JJ-OX-004 and compound 3 (Fig. 6, G and H, and Supplementary Data 5). Both compounds share the same oxadiazolone warhead and an aromatic ring but differ in their target selectivity. JJ-OX-004 is a selective inhibitor of FphE with only minor cross reactivity with FphB in *staphylococcus* bacteria, whereas compound 3 is a more general antimicrobial agent with other targets including FabH which is not a serine hydrolase (*14, 23*). Interactions near the active site, including the GSSSG motif at the catalytic serine, are similar and highly conserved (Fig. 6G; Gly27, Ala28 and Pro129). The unique buried pocket occupied by compound 3 is formed by residues that are also highly conserved (Fig. 6, G and H; Asn29, Ser102, Asn144, Met160, Phe163, Ile169, Met177, Trp197 and Tyr260). In contrast, the tail of JJ-OX-004 does not enter this pocket but reaches the surface via a hydrophobic opening that is formed by the side chain of moderately conserved Leu156 and the backbone of less conserved residues (Fig. 6, G and H; Glu157, Ser178 and Tyr191). JB101 occupies the same distal hydrophobic tunnel as JJ-OX-004 but bears a fluorophosphonate (FP) warhead, which has much higher reactivity and thus broader serine hydrolase labeling properties compared to the oxadiazolones (*57*). This increased reactivity likely explains the reduced target selectivity of the FP probe despite the similar pocket engagement profile to JJ-OX-004. These observations indicate that both the location of binding within FphE and the intrinsic reactivity profile of the warhead contribute to selectivity outcomes. Our findings suggest that the presence of an optimal electrophilic warhead, an aromatic ring, and a hydrophobic pocket-occupying substituent produce molecules with a high degree of selectivity for FphE.

## Discussion

Although serine hydrolases play crucial roles in bacterial physiology and pathogenesis, systemic characterization of this family of enzymes in *S. aureus* has only recently been initiated. The discovery of the Fph family has provided new insights into biofilm formation and virulence in *S. aureus* (*11, 17, 18*), as well as identified new targets for the development of species-specific molecular imaging tools (*14*). Among this serine hydrolase family, FphE stands out as a structurally unique and pathogen-selective target. Our findings presented here provide an integrated structural, biochemical, and bioinformatic characterization of FphE uncovering a cross-monomer homodimer configuration that is unique for a serine hydrolase. This work also provides a structural and bioinformatic basis for future rational design of imaging probes or inhibitors targeting this enzyme.

Unlike most members of the Fph family, FphE exhibits a restricted distribution within *Staphylococcus* species and is absent in the representative nosocomial coagulase-negative staphylococci (CoNS), *Staphylococcus epidermidis* and *Staphylococcus hemolyticus* (*10, 14, 58*). This taxonomic exclusivity positions it as a potential selective biomarker for *S. aureus* infections. The ability to discriminate *S. aureus* from other related bacterial infections is critical in clinical settings where rapid and efficient identification of a specific bacteria affects treatment choices. Beyond the diagnostic potential of FphE, its exact functional role remains unclear.

Broad variability in *fphE* transcriptional responses to external stimuli (*17, 25–37*) suggest that it may participate in adaptive processes that enable *S. aureus* to respond to environmental and host-associated conditions, rather than serving a single dedicated metabolic function inside the bacteria. Transcriptomic analyses indicate that *fphE* (SA2367, SACOL2597) expression is controlled by the alternative sigma factor, σ^B^ regulon, with expression patterns varying across stress conditions (*25–27*). *fphE* induction has been observed in response to nitroxyl donor Angeli’s salt (NWMN_2480 and SA2367) (*28*), mupirocin (SACOL2597) (*29*), tea tree oil (SACOL2597) (*30*), and rhodomyrtone (SAR2661) (*31*), while repression occurs with diclofenac (SACOL2597) (*27*), *menD* mutation (SACOL2597) (*59*), or by interspecies competition (SAUSA300_2518) (*32*). Expression is further modulated across growth states (strain UAMS-1) (*34*) and within vancomycin-intermediate resistant (strain SA137/93G) and susceptible (strain SA1450/94) strains (*33*), suggesting its role in adaptive responses. RST4–1 represents a major human genotype of *S. aureus*, but its occasional isolation from bovine mastitis cases raised the possibility of pseudogenization of *fphE* (*60*) that suggests it may contribute to host-specific adaptation. Functional studies provide additional evidence that FphE contributes to the adaptive response, as *fphE* disruption reduced human endothelial cell damage (*35*), and we reported that *fphE* transposon mutant displayed reduced liver bacterial load without affecting heart or kidney colonization (*17*). Several proteomic studies (*61–66*) have confirmed the presence of the FphE protein in various *S. aureus* strains and growth conditions without directly linking FphE function to a specific phenotype but giving further support to the hypothesis that FphE has a general role in adaptive responses to environmental stresses. Notebly, one study showed that FphE could hydrolyze nitrophenyl acetate, demonstrating esterase activity toward an aromatic ester (*66*).

Our previous study demonstrated that JJ-OX-004, which bears an aromatic ring on the scaffold, exhibited greater selectivity for the active site serine on FphE than JJ-OX-003 (*14*). While the initial probe design was guided by an existing scaffold that contains both an aromatic ring and an oxadiazolone electrophile rather than by direct structural information about the target (*23*), adding the benzene ring to the JJ-OX-003 scaffold proved essential for labeling FphE. In our present study, we use comparative genomic and structural analyses to provide a mechanistic rationale for the observed preference of FphE for scaffolds that contain an aromatic ring adjacent to the reactive electrophile. We found that *fphE* is co-localized on a genomic locus with aromatic compound-processing amidohydrolases (SAUSA300_2517 and its homologs), suggesting that the natural substrate of FphE may be aromatic in nature. Nevertheless, our results collectively highlight how integrating bioinformatic analysis such as gene colocalization and phylogenetic analysis with chemical design strategies can guide the development of more potent and selective activity-based probes.

Our biochemical analysis of recombinant FphE suggests that this enzyme forms a stable dimer that resists dissociation even when diluted. This robustness of FphE dimerization suggests a functional or structural necessity of the cross-monomeric assembly. The FphE fluorescent probe JJ-OX-007 efficiently labels both monomeric and dimeric forms without inducing interconversion, confirming that catalytic accessibility is maintained in the dimeric state. Also, this result suggests that JJ-OX-007 minimally perturbs the oligomeric equilibrium of FphE and may serve as a valuable tool for the biochemical study of FphE. However, the higher hydrolytic activity of the dimer toward the 4-MUC substrate suggests subtle structural differences in the active site pocket that enhance catalytic efficiency in the dimeric conformation. Interestingly, co-incubation with the substrate 4-MUC significantly slowed the dimer-to-monomer conversion at 37 °C. This observation implies active-site engagement may stabilize the dimeric assembly. Temperature dependence of the dimeric status further suggests that dynamic equilibrium may be influenced by environmental factors, which may have physiological relevance under stress conditions where oligomeric transitions could modulate enzyme activity.

Our results indicate that there are general alpha/beta hydrolase homologs of the FphE monomer in the PDB but none that are close enough matches to provide structure-function information. While FphE is a member of a family of enzymes with a conserved and well-characterized fold, the homodimer we describe here appears to have never been reported for any other alpha/beta hydrolase members and the dimer architecture is not easily predicted using current modeling methods. SAXS data confirms that the observed crystal architecture is highly similar to the in-solution confirmation and clearly differs from other alpha/beta hydrolase homodimer interface, as seen for related hydrolases such as FphF. The FphE homodimer crystal forms suggest a level of flexibility of the overall fold that is not dependent on a traditional lid arrangement, presenting another unique feature of this architecture. All these aspects have implications for the binding of substrates or inhibitors.

To our knowledge, the structure of JB101 bound in the active site of FphE is the first FP bound structure within the *S. aureus* FphA-J proteins, even though an FP probe was used to identify this class of enzymes. Our structure can be used to describe FphA-J protein interactions with FPs in general. Such FP-alkynes are used frequently to target active site serine residues and a crystal structure of a computationally designed serine hydrolase with the FP-ligand covalently bound has been reported (*67*). Several other studies have also reported on FP ligands that are covalently bound to serine active site residues, including FAS thioesterase (*68*) and phospholipases A (*69*). The structure of JB101 bound in the active site of FphE, when combined with the JJ-OX-004 structure (*14*) offers a framework for further structure-guided probe design and optimization.

The compound 3 bound structure, on the other hand, reveals an alternative interaction site within the unique homodimer architecture. This molecule binds such that it occupies multiple binding regions in and near the active site. This includes the active site triad, which is bound in manner similar to all other covalent bound ligands for this class of hydrolases (fig. S13). In addition, the first phenyl ring of compound 3 occupies a space pointing towards the protein surface, similar to JB101, JJ-OX-004 (*14*) and a boronic acid ligand Z27, previously described by us (fig. S13, A to D) (*18*). Lastly, the final two phenyl rings bend away from the surface to occupy a space only observed for compound 3, suggesting a potentially high level of plasticity in inhibitor/substrate binding.

The high conservation of residues that interact with the bound inhibitors provides a structural rationale for the contrasting selectivity of JJ-OX-004 and compound 3, which share the same oxadiazolone electrophile and adjacent aromatic ring but differ in the substituent attached via an amide linkage. Compound 3 interacts with highly conserved regions of the active site and a neighboring pocket, whereas JJ-OX-004 engages less conserved peripheral residues within a hydrophobic channel unique to FphE. This interaction networks not only partially explains the selectivity of JJ-OX-004 for FphE but also provides a generalizable design principle for targeting non-conserved hydrophobic pockets outside the catalytic triad to achieve FphE specificity. These insights show that combining structural data with conservation analysis from bioinformatics can effectively guide future rational design of selective probes and inhibitors. Taken together, these findings advance our understanding of FphE from a poorly characterized serine hydrolase to a structurally distinct enzyme with a potentially conserved metabolic role. It also establishes a rationale for further development of potential *S. aureus* specific probes and inhibitors.

## Materials and methods

### Bioinformatic analysis

To identify FphE homologs, PSI-BLAST searches were performed against the NCBI non-redundant protein database downloaded in February 2024 using FphE as a query, a cut-off E-value of 1e-50, and a minimum coverage of 0.70 for 7 iterations (*70*). Redundant sequences were removed by clustering 11,030 proteins at 95 % identity and 95% coverage with MMseqs2 (version 16.747c6), yielding 6,141 representative sequences (*71*). Sequences were aligned using MAFFT v7.525 (--globalpair --maxiterate 1000) and the resulting alignment was used to generate a phylogenetic by FastTree 2 (version 2.1.11) (*72, 73*). Genomic loci and taxonomic classifications were retrieved from NCBI nucleotide and taxonomy databases. Protein domains were annotated using HMMER 3.4 (*74*) and MMseqs2 (*71*). The tree was visualized with the R package ggtree v3.10.1 (*75*). Structural models were generated by AlphaFold3 (*53*) or downloaded from AlphaFold database (*76*), and visualized by ChimeraX 1.10 (*77*). To identify structural homologs, Foldseek easy-search (version d7a5a0ef39642aab76b63dcac0aafdf098f623e9) was performed against the PDB database using the AlphaFold structure of SAUSA300_2517 as a query, a cut-off E-value of 1e-20, and a minimum coverage of 0.80 (*78*). Sequence logos were created using WebLogo (*56*).

### Chemicals

Q41 (CAS number 1430628-65-1) was purchased from Enamine Ltd. Oxadiazolone probes compound 3 and JJ-OX-007 were synthesized as previously reported (*14, 23*). The synthesis scheme and detailed protocols for JB101 along with NMR spectra are provided in the Supplementary Information.

### FphE expression, purification and crystallization

Full-length FphE from *S. aureus* USA300 (Uniprot ID Q2FDS6, SAUSA300_2518) was expressed and purified as previously described via a pET-28a based *Escherichia coli* expression system (*14*). The N-terminal His6-tag was always cleaved off using a 3C protease for structural studies. The final purification step was in all cases size exclusion with the used buffer indicated in brackets in the following paragraphs. The final step prior to crystallization or incubation with ligands was concentrating using centrifugal concentrators with the final concentration in mg/mL indicated for each crystallization setup. Crystals were always grown at 16 °C.

For JB101 bound in crystal form 1, 20 µL 25 mg/mL FphE (10 mM HEPES pH 7.5, 100 mM NaCl) were mixed with 3 µL JB101 (10 mM in DMSO) and incubated at 4 °C overnight. 0.2 µL FphE-JB101 were mixed with 0.2 µL of reservoir solution. Sitting drop reservoir contained 25 µL of 180 mM magnesium chloride hexahydrate, 100 mM Tris pH 8.0, 22.5 % PEG 2000 MME. Crystal was frozen in a solution of ∼25 % glycerol, 75 % reservoir. For compound 3 bound in crystal form 2, 65 µL 16 mg/mL FphE (10 mM HEPES pH 7.6, 100 mM NaCl) were mixed with 7.5 µL compound 3 (10 mM in DMSO) and incubated at 4 °C overnight. 0.15 µL FphE-compound-3 solution was mixed with 0.3 µL of reservoir solution. Sitting drop reservoir contained 25 µL of 160 mM potassium thiocyanate, 180 mM Tris pH 8.5, 20 % PEG 2000 MME. Crystal was frozen in a solution of ∼25 % ethylene glycol, 75 % reservoir. For unbound crystal form 3, 0.3 µL 14 mg/mL FphE (10 mM HEPES pH 7.6, 100 mM NaCl) were mixed with 0.3 µL of reservoir solution. Sitting drop reservoir contained 25 µL of 200 mM magnesium chloride hexahydrate, 100 mM Tris pH 8.5, 25 % PEG 2000 MME. Crystal was frozen in a solution of ∼25 % glycerol, 75 % reservoir. For unbound crystal form 5, 0.3 µL 14 mg/mL FphE (10 mM HEPES pH 7.6, 100 mM NaCl) were mixed with 0.15 µL of reservoir solution. Sitting drop reservoir contained 25 µL of 200 mM potassium thiocyanate, 100 mM Tris pH 8.5 and 22 % w/v PEG 2,000 MME. Crystal was frozen in a solution of ∼25 % glycerol, 75 % reservoir. For Q41 bound in crystal form 6, 16 µL 11 mg/mL FphE (10 mM HEPES pH 7.5, 100 mM NaCl) were mixed with 6 µL Q41 (50 mM in DMSO) and incubated at 18 °C overnight. 0.15 µL FphE-Q41 solution was mixed with 0.3 µL of reservoir solution. Sitting drop reservoir contained 25 µL of 180 mM potassium thiocyanate, 100 mM sodium acetate pH 5.5, 22.5 % PEG 2000 MME. Crystal was frozen in a solution of ∼25 % glycerol, 75 % reservoir. For unbound crystal form 8, 0.3 µL 11 mg/mL FphE (10 mM HEPES pH 7.5, 100 mM NaCl) were mixed with 0.15 µL of reservoir solution. Sitting drop reservoir contained 25 µL of 180 mM potassium thiocyanate, 100 mM Tris pH 8.5 and 22.5 % w/v PEG 2,000 MME. Crystal was frozen in a solution of ∼25 % glycerol, 75 % reservoir. Crystallization information about all FphE crystal forms, as well as not fully processed and untested FphE crystals are listed in tables S4, S5 and S6.

### Analysis of oligomeric status and probe labeling of FphE monomer and dimer

Purified recombinant FphE monomer (peak 2) and dimer (peak 1) fractions were aliquoted as 2.0 µM in 1X PBS solution (20 µL per each sample) and incubated with or without 4-methylumbelliferyl caprylate (4-MUC, 500 µM final concentration; 1 µL of 10 mM stock in DMSO) for 30 min at ambient temperature. After the preincubation, probe JJ-OX-007 (1 µM final concentration; 0.1% DMSO stock in 1X PBS, v/v) was added, and the mixtures were further incubated at 37 °C for 1 h. As controls, samples containing DMSO vehicle were also prepared. After incubation, each sample was mixed with 20 µL of non-reducing, non-denaturing 4X native-PAGE loading buffer and immediately subjected to gel electrophoresis without boiling. A total of 300 ng of protein was loaded per lane onto a 12% native-polyacrylamide gel and run at 100 V under ambient temperature. Fluorescently labeled proteins were visualized using a GE Typhoon FLA 9000 (GE Healthcare, Pittsburgh, PA).

### Analysis of FphE dimeric stability and probe labeling under variable temperatures

To a purified dimeric FphE (2.0 µM, peak 1) in 20 µL of 1X PBS solution, probe JJ-OX-007 (1 µM final concentration; 0.1% DMSO stock in 1X PBS, v/v) or DMSO carrier (0.1% DMSO in 1X PBS, v/v) was added in the presence or absence of 4-methylumbelliferyl caprylate (4-MUC, 500 µM final concentration; 1 µL of 10 mM stock in DMSO). After 1 h incubation under the indicated temperatures (4 °C, room temperature, or 37 °C), each sample was mixed with 20 µL of non-reducing, non-denaturing 4X native-PAGE loading buffer and immediately subjected to gel electrophoresis without boiling. A total of 300 ng of protein was loaded per lane onto a 12% native-polyacrylamide gel and run at 100 V under ambient temperature. Fluorescently labeled proteins were visualized using a GE Typhoon FLA 9000 (GE Healthcare, Pittsburgh, PA). After scanning, the gels were stained with Coomassie Brilliant Blue to visualize total protein content.

### Monomer/dimer ratio quantification on native gels

After scanning, the gels were stained with Coomassie Brilliant Blue to visualize total protein content. Band intensities were quantified using the software ImageJ from native-PAGE gels. Images were converted to 8-bit grayscale and background was subtracted. Rectangular ROIs of identical size were placed over the hands in each lane, and integrated density (area under the peak) was recorded. Because no loading control was used, per-lane normalization was applied: Monomer fraction = *I*_M_/(*I*_M_ + *I*_D_), Dimer fraction = *I*_D_/(*I*_M_ + *I*_D_), where *I*_M_ and *I*_D_ are the integrated densities of the monomer and dimer bands, respectively. Only lanes from the same gel and exposure were compared.

### Enzymatic assay

The purified dimeric and monomeric FphE (0.5 nM) in 10 µL of 1X PBS solution, was added in the presence or absence of varied concentration of 4-methylumbelliferyl caprylate (4-MUC, 0–25 µM final concentration and fluorescence (λ_ex_ = 365 nm and λ_em_ = 455 nm) was measured at 30 °C in 1 min intervals on a Cytation 3 imaging reader (BioTek, Winooski, VT, USA) for 60 min. Turnover rates in the linear phase of the reaction (10–20 min) were calculated using GraphPad prism 10.0 software as RFU/min. Rates were normalized by subtracting background hydrolysis rates measured for each substrate concentrations in reaction buffer in the absence of protein.

### FphE crystallography data collection, processing, refinement, deposition, and analysis

Data was collected at the Australian synchrotron MX1 and MX2 beamlines (*79, 80*). Data sets were processed with XDS (*81*), merging and scaling were performed using AIMLESS (*82*). Phases were solved with Phenix Phaser molecular replacement (*83*) using various FphE structures from different crystal forms. In many cases phaser solutions were placed incorrectly with the dimer helices stretching into a neighboring crystal molecule. Only searches using dimers often lacking the connecting helix or the use of chain segments, most frequently 1 to 151 and 172 to 276, resulted in the correct solution. Model building and refinement were conducted in COOT (*84*) and Phenix (*85*). The final structure was deposited to the worldwide protein databank (PDB) (*47*) JB101 bound in crystal form 1 at 1.49 Å (8SBQ), Compound-3 bound in crystal form 2 at 1.60 Å (8G49), unbound crystal form 3 at 1.95 Å (8G48), unbound crystal form 5 at 2.04 Å (9COM), Q41 bound in crystal form 6 at 2.50 Å (9EBF), unbound crystal form 8 at 2.30 Å (9D87) and unbound crystal form 9 at 2.60 Å (9EDJ). Statistics for the data sets are listed in tables S7 and S8. Structure figures, analysis and alignments were created with UCSF Chimera (*86*), ChimeraX (*77*), and LigPlot+ (*87*). 3D protein structure similarity search was performed using the DALI (*88*) Web server (table S11, S12 and S13).

### Small-Angle X-ray Scattering (SAXS) data collection, processing, and analysis

Full-length FphF from *S. aureus* USA300 (Uniprot ID A0A0H2XFN6, SAUSA300_2564) was purified as previously described in detail (*9*). FphE and FphF SAXS experiments were performed at the Australian Synchrotron, ANSTO BioSAXS beamline and followed a procedure previously described for FphH (*15*). Collection parameters are summarized in table S10. An in-line size exclusion chromatography (SEC) SAXS mode (*89*) was used on the CoFlow Autoloader fit with an autosampler and a Knauer HPLC Pump. 50 µL of either FphE (1.0 mg/mL) or FphF (6.5 mg/mL) in 10 mM HEPES pH 7.5, 50 mM NaCl were injected onto a Superdex Increase S75 5/150 GL column (Cytiva). The size exclusion contained 10 mM HEPES pH 7.5, 50 mM NaCl and 0.1% sodium azide, with the SAXS capillary as the sheath fluid (*90*) containing the same buffer. Data calibration and reduction were also performed as previously described for FphH (*15*). Reduced data were processed and analyzed using the ATSAS suite of software (*91*). Chromixs (*92*) was used for buffer subtraction. Structural parameters shown in table S7 were calculated using the RAW (*93*) interface, and the particle distribution function was calculated using GNOM (*94*) Ab-initio shape determination was done in DAMMIF (*95*) and DAMAVER (*96*). The FphE dimer crystal structure (PDB ID 8T87) or all possible dimer combinations in the FphF tetramer crystal structure (PDB ID 6VH9) were fitted to the experimental data using Crysol (*91*). SREFLEX (*97*) was used to improve model agreement with the SAXS data. The FphE SAXS dataset was deposited to the SASBDB under ID SASDX64, FphF under ID SASDX74.

### Protease substrate analysis

FphE was screened for protease activity against eight commercially available fluorogenic peptide model substrates as previously described for FphH (*11*) and FphI (*15*) (table S2).

## Acknowledgments

We would like to thank Alexander T. Bakker, Nathaniel I. Martin and Mario van der Stelt from Leiden University for providing compound 3. We thank Dr. Anthony O’Donoghue from University of California, San Diego for performing the protease activity experiment. We also thank Dr. Yuri Choi at Stanford University for helpful discussions. Molecular graphics and analyses performed with UCSF ChimeraX, developed by the Resource for Biocomputing, Visualization, and Informatics at the University of California, San Francisco, with support from National Institutes of Health R01-GM129325 and the Office of Cyber Infrastructure and Computational Biology, National Institute of Allergy and Infectious Diseases. This research was undertaken in part using the MX1 and MX2 beamlines at the Australian Synchrotron, part of ANSTO, and made use of the Australian Cancer Research Foundation (ACRF) detector.

## Funding

New Zealand Marsden Fund Council from government funding, managed by Royal Society Te Apa̅rangi (MF)

The Sir Charles Hercus Health Research Fellowship of the Health Research Council of New Zealand (MF)

National Institutes of Health R01 EB026332 (MB)

## Author contributions

Conceptualization: MF, JJ, MB

Methodology: MF, JJ, XU, TU, JMB, HL

Investigation: MF, JJ, XU, TU, JMB, HL

Visualization: MF, XU, JJ

Formal analysis: MF, JJ, TU, HL

Funding acquisition: MF, MB

Project administration: MF, JJ, MB

Supervision: MF, MB

Writing: MF, JJ, MB

## Data and materials availability

All data needed to evaluate the conclusions in the paper are present in the paper and/or the Supplementary Materials. The crystal structure datasets generated and analyzed during the current study are available in the worldwide Protein Data Bank under PDB IDs 8SBQ, 8G49, 8G48, 9COM, 9EBF, 9D87 and 9EDJ. Authors will release the atomic coordinates and experimental data upon article publication. SAXS datasets are available in the Small Angle Scattering Biological Data Bank SASBDB IDs SASDX64 and SASDX74, they will be released upon article publication.

## Supplementary Materials

### This PDF file includes

Figs. S1 to S13

Tables S1 to S13

Chemical synthesis and NMR spectra

References

### Other Supplementary Material for this manuscript includes the following

Supplementary Data 1 to 5

## References and Notes

1. H. F. Wertheim et al., The role of nasal carriage in *Staphylococcus aureus* infections. Lancet Infect Dis 5, 751–762 (2005).

2. S. Y. Tong, J. S. Davis, E. Eichenberger, T. L. Holland, V. G. Fowler, Jr., *Staphylococcus aureus* infections: epidemiology, pathophysiology, clinical manifestations, and management. Clin Microbiol Rev 28, 603–661 (2015).

3. K. B. Laupland et al., The changing epidemiology of *Staphylococcus aureus* bloodstream infection: a multinational population-based surveillance study. Clin Microbiol Infect 19, 465–471 (2013).

4. C. de la Fuente-Nunez, F. Reffuveille, L. Fernandez, R. E. W. Hancock, Bacterial biofilm development as a multicellular adaptation: antibiotic resistance and new therapeutic strategies. Curr Opin Microbiol 16, 580–589 (2013).

5. S. Y. C. Tong, J. S. Davis, E. Eichenberger, T. L. Holland, V. G. Fowler, *Staphylococcus aureus* infections: epidemiology, pathophysiology, clinical manifestations, and management. Clin Microbiol Rev 28, 603–661 (2015).

6. M. Clauss, U. F. Tafin, A. Bizzini, A. Trampuz, T. Ilchmann, Biofilm formation by staphylococci on fresh, fresh-frozen and processed human and bovine bone grafts. Eur Cell Mater 25, 159–166 (2013).

7. J. Z. Long, B. F. Cravatt, The Metabolic Serine Hydrolases and Their Functions in Mammalian Physiology and Disease. Chem Rev 111, 6022–6063 (2011).

8. L. Chen, L. J. Keller, E. Cordasco, M. Bogyo, C. S. Lentz, Fluorescent triazole urea activity-based probes for the single-cell phenotypic characterization of *Staphylococcus aureus*. Angew Chem Int Ed Engl 58, 5643–5647 (2019).

9. M. Fellner et al., Structural Basis for the Inhibitor and Substrate Specificity of the Unique Fph Serine Hydrolases of Staphylococcus aureus. ACS Infect Dis 6, 2771–2782 (2020).

10. L. J. Keller et al., Characterization of Serine Hydrolases Across Clinical Isolates of Commensal Skin Bacteria Staphylococcus epidermidis Using Activity-Based Protein Profiling. ACS Infect Dis 6, 930–938 (2020).

11. M. Fellner et al., Biochemical and Cellular Characterization of the Function of Fluorophosphonate-Binding Hydrolase H (FphH) in Staphylococcus aureus Support a Role in Bacterial Stress Response. ACS Infect Dis 9, 2119–2132 (2023).

12. S. Chen et al., Identification of highly selective covalent inhibitors by phage display. Nat Biotechnol 39, 490–498 (2021).

13. M. Fellner, Newly discovered Staphylococcus aureus serine hydrolase probe and drug targets. ADMET DMPK 10, 107–114 (2022).

14. J. Jo et al., Development of Oxadiazolone Activity-Based Probes Targeting FphE for Specific Detection of Staphylococcus aureus Infections. J Am Chem Soc 146, 6880–6892 (2024).

15. M. Fellner et al., Similar but Distinct-Biochemical Characterization of the Staphylococcus aureus Serine Hydrolases FphH and FphI. Proteins, (2024).

16. S. Wang et al., An mRNA Display Approach for Covalent Targeting of a Staphylococcus aureus Virulence Factor. Journal of the American Chemical Society 147, 8312–8325 (2025).

17. C. S. Lentz et al., Identification of a *S. aureus* virulence factor by activity-based protein profiling (ABPP). Nat Chem Biol 14, 609–617 (2018).

18. T. Upadhyay et al., Identification of covalent inhibitors of Staphylococcus aureus serine hydrolases important for virulence and biofilm formation. Nature Communications 16, 5046 (2025).

19. G. M. Simon, B. F. Cravatt, Activity-based Proteomics of Enzyme Superfamilies: Serine Hydrolases as a Case Study. Journal of Biological Chemistry 285, 11051–11055 (2010).

20. J. D. Schrag, M. Cygler, Lipases and alpha/beta hydrolase fold. Methods Enzymol 284, 85–107 (1997).

21. I. Staub, S. A. Sieber, beta-Lactam Probes As Selective Chemical-Proteomic Tools for the Identification and Functional Characterization of Resistance Associated Enzymes in MRSA. Journal of the American Chemical Society 131, 6271–6276 (2009).

22. M. J. Uddin, H. S. Overkleeft, C. Lentz, Activity-Based Protein Profiling in Methicillin-Resistant Staphylococcus aureus Reveals Broad Reactivity of a Carmofur-Derived Probe. Chembiochem, e202300473 (2023).

23. A. T. Bakker et al., Chemical Proteomics Reveals Antibiotic Targets of Oxadiazolones in MRSA. J Am Chem Soc 145, 1136–1143 (2023).

24. K. S. Ikuta et al., Global mortality associated with 33 bacterial pathogens in 2019: a systematic analysis for the Global Burden of Disease Study 2019. The Lancet 400, 2221–2248 (2022).

25. M. Bischoff et al., Microarray-based analysis of the σ regulon. Journal of Bacteriology 186, 4085–4099 (2004).

26. S. Fuchs et al., Aureolib - a proteome signature library: towards an understanding of staphylococcus aureus pathophysiology. PLoS One 8, e70669 (2013).

27. J. T. Riordan et al., Alterations in the transcriptome and antibiotic susceptibility of Staphylococcus aureus grown in the presence of diclofenac. Ann Clin Microbiol Antimicrob 10, 30 (2011).

28. H. Peng et al., Sulfide Homeostasis and Nitroxyl Intersect via Formation of Reactive Sulfur Species in Staphylococcus aureus. mSphere 2, (2017).

29. S. Reiss et al., Global analysis of the Staphylococcus aureus response to mupirocin. Antimicrob Agents Chemother 56, 787–804 (2012).

30. J. A. Cuaron et al., Tea tree oil-induced transcriptional alterations in Staphylococcus aureus. Phytother Res 27, 390–396 (2013).

31. W. Sianglum, P. Srimanote, P. W. Taylor, H. Rosado, S. P. Voravuthikunchai, Transcriptome analysis of responses to rhodomyrtone in methicillin-resistant Staphylococcus aureus. PLoS One 7, e45744 (2012).

32. M. Wei, et al., An exploration of mechanisms underlying Desemzia incerta colonization resistance to methicillin-resistant Staphylococcus aureus on the skin. Msphere 9, e00636–00623 (2024).

33. A. Jansen et al., Production of capsular polysaccharide does not influence vancomycin susceptibility. Bmc Microbiology 13, (2013).

34. K. E. Beenken et al., Global gene expression in Staphylococcus aureus biofilms. J Bacteriol 186, 4665–4684 (2004).

35. X. Xiao, Y. Li, L. Li, Y. Q. Xiong, Identification of Methicillin-Resistant Staphylococcus aureus (MRSA) Genetic Factors Involved in Human Endothelial Cells Damage, an Important Phenotype Correlated with Persistent Endovascular Infection. Antibiotics 11, 316 (2022).

36. N. Islam, J. M. Ross, M. R. Marten, Proteome Analyses of Staphylococcus aureus Biofilm at Elevated Levels of NaCl. Clin Microbiol 4, (2015).

37. C. J. Frapwell et al., Antimicrobial Activity of the Quinoline Derivative HT61 against Staphylococcus aureus Biofilms. Antimicrob Agents Chemother 64, (2020).

38. M. Begley, R. D. Sleator, C. G. Gahan, C. Hill, Contribution of three bile-associated loci, bsh, pva, and btlB, to gastrointestinal persistence and bile tolerance of Listeria monocytogenes. Infect Immun 73, 894–904 (2005).

39. A. K. Marr et al., Overexpression of PrfA leads to growth inhibition of in glucose-containing culture media by interfering with glucose uptake. Journal of Bacteriology 188, 3887–3901 (2006).

40. L. O. Henderson et al., Transcriptional profiling of the L. monocytogenes PrfA regulon identifies six novel putative PrfA-regulated genes. FEMS Microbiol Lett 367, (2020).

41. C. Becavin et al., Listeriomics: an Interactive Web Platform for Systems Biology of Listeria. mSystems 2, (2017).

42. A. Ray, K. A. Edmonds, L. D. Palmer, E. P. Skaar, D. P. Giedroc, Glucose-Induced Biofilm Accessory Protein A (GbaA) Is a Monothiol-Dependent Electrophile Sensor. Biochemistry 59, 2882–2895 (2020).

43. L. Yu et al., A novel repressor of the ica locus discovered in clinically isolated super-biofilm-elaborating Staphylococcus aureus. MBio 8, 10.1128/mbio.02282-02216 (2017).

44. Y. Fan et al., The catalytic mechanism of direction-dependent interactions for 2, 3-dihydroxybenzoate decarboxylase. Applied Microbiology and Biotechnology 107, 7451–7462 (2023).

45. X. Gao et al., Structural basis of salicylic acid decarboxylase reveals a unique substrate recognition mode and access channel. Journal of Agricultural and Food Chemistry 69, 11616–11625 (2021).

46. M. Song et al., 2, 3-Dihydroxybenzoic Acid Decarboxylase from Fusarium oxysporum: Crystal Structures and Substrate Recognition Mechanism. ChemBioChem 21, 2950–2956 (2020).

47. H. Berman, K. Henrick, H. Nakamura, J. L. Markley, The worldwide Protein Data Bank (wwPDB): ensuring a single, uniform archive of PDB data. Nucleic Acids Res 35, D301–303 (2007).

48. K. E. Jaeger, B. W. Dijkstra, M. T. Reetz, Bacterial biocatalysts: molecular biology, three-dimensional structures, and biotechnological applications of lipases. Annu Rev Microbiol 53, 315–351 (1999).

49. J. Jumper et al., Highly accurate protein structure prediction with AlphaFold. Nature 596, 583–589 (2021).

50. T. Paysan-Lafosse et al., InterPro in 2022. Nucleic Acids Res 51, D418–D427 (2023).

51. L. Holm, A. Laiho, P. Toronen, M. Salgado, DALI shines a light on remote homologs: One hundred discoveries. Protein Sci 32, e4519 (2023).

52. Q. Qi et al., The structure of a complex of the lactonohydrolase zearalenone hydrolase with the hydrolysis product of zearalenone at 1.60 Å resolution. Acta Crystallogr F 73, 376–381 (2017).

53. J. Abramson et al., Accurate structure prediction of biomolecular interactions with AlphaFold 3. Nature, (2024).

54. D. L. Ollis et al., The alpha/beta hydrolase fold. Protein Eng 5, 197–211 (1992).

55. M. Nardini, D. A. Lang, K. Liebeton, K. E. Jaeger, B. W. Dijkstra, Crystal structure of pseudomonas aeruginosa lipase in the open conformation. The prototype for family I.1 of bacterial lipases. J Biol Chem 275, 31219–31225 (2000).

56. G. E. Crooks, G. Hon, J.-M. Chandonia, S. E. Brenner, WebLogo: a sequence logo generator. Genome research 14, 1188–1190 (2004).

57. F. Faucher, J. M. Bennett, M. Bogyo, S. Lovell, Strategies for tuning the selectivity of chemical probes that target serine hydrolases. Cell chemical biology 27, 937–952 (2020).

58. L. J. Keller, M. Lakemeyer, M. Bogyo, in Methods in Enzymology. (Elsevier, 2022), vol. 664, pp. 1–22.

59. C. Kohler et al., A defect in menadione biosynthesis induces global changes in gene expression in Staphylococcus aureus. J Bacteriol 190, 6351–6364 (2008).

60. D. S. Ko et al., Comparative genomics of bovine mastitis-origin Staphylococcus aureus strains classified into prevalent human genotypes. Res Vet Sci 139, 67–77 (2021).

61. C. J. Caballero et al., The regulon of the RNA chaperone CspA and its auto-regulation in Staphylococcus aureus. Nucleic Acids Res 46, 1345–1361 (2018).

62. C. Kohler, et al., Proteomic and Membrane Lipid Correlates of Re-duced Host Defense Peptide Susceptibility in a snoD Mutant of Staphylococcus aureus. Antibiotics-Basel 8, (2019).

63. D. Frees et al., New Insights into Stress Tolerance and Virulence Regulation from an Analysis of the Role of the ClpP Protease in the Strains Newman, COL, and SA564. Journal of Proteome Research 11, 95–108 (2012).

64. D. Becher et al., A proteomic view of an important human pathogen--towards the quantification of the entire Staphylococcus aureus proteome. PLoS One 4, e8176 (2009).

65. S. Michalik et al., A global Staphylococcus aureus proteome resource applied to the in vivo characterization of host-pathogen interactions. Sci Rep 7, 9718 (2017).

66. B. D. Saylor, J. J. Love, A secreted Staphylococcus aureus lipase engineered for enhanced alcohol affinity for fatty acid esterification. Journal of Molecular Catalysis B: Enzymatic 133, S44–S52 (2016).

67. S. Rajagopalan et al., Design of activated serine-containing catalytic triads with atomic-level accuracy. Nat Chem Biol 10, 386–391 (2014).

68. W. Zhang et al., Crystal structure of FAS thioesterase domain with polyunsaturated fatty acyl adduct and inhibition by dihomo-gamma-linolenic acid. Proc Natl Acad Sci U S A 108, 15757–15762 (2011).

69. H. Wang et al., Structure of Human GIVD Cytosolic Phospholipase AReveals Insights into Substrate Recognition. Journal of Molecular Biology 428, 2769–2779 (2016).

70. S. F. Altschul et al., Gapped BLAST and PSI-BLAST: a new generation of protein database search programs. Nucleic acids research 25, 3389–3402 (1997).

71. M. Steinegger, J. Söding, MMseqs2 enables sensitive protein sequence searching for the analysis of massive data sets. Nature biotechnology 35, 1026–1028 (2017).

72. K. Katoh, D. M. Standley, MAFFT multiple sequence alignment software version 7: improvements in performance and usability. Molecular biology and evolution 30, 772–780 (2013).

73. M. N. Price, P. S. Dehal, A. P. Arkin, FastTree 2–approximately maximum-likelihood trees for large alignments. PloS one 5, e9490 (2010).

74. S. R. Eddy, in Genome Informatics 2009: Genome Informatics Series Vol. 23. (World Scientific, 2009), pp. 205–211.

75. S. Xu, et al., Ggtree: a serialized data object for visualization of a phylogenetic tree and annotation data. Imeta 1, e56 (2022).

76. M. Varadi et al., AlphaFold Protein Structure Database: massively expanding the structural coverage of protein-sequence space with high-accuracy models. Nucleic acids research 50, D439–D444 (2022).

77. E. C. Meng et al., UCSF ChimeraX: Tools for structure building and analysis. Protein Sci 32, e4792 (2023).

78. M. Van Kempen et al., Fast and accurate protein structure search with Foldseek. Nature biotechnology 42, 243–246 (2024).

79. D. Aragao et al., MX2: a high-flux undulator microfocus beamline serving both the chemical and macromolecular crystallography communities at the Australian Synchrotron. J Synchrotron Radiat 25, 885–891 (2018).

80. N. P. Cowieson et al., MX1: a bending-magnet crystallography beamline serving both chemical and macromolecular crystallography communities at the Australian Synchrotron. J Synchrotron Radiat 22, 187–190 (2015).

81. W. Kabsch, Xds. Acta Crystallogr D Biol Crystallogr 66, 125–132 (2010).

82. M. D. Winn et al., Overview of the CCP4 suite and current developments. Acta Crystallogr D Biol Crystallogr 67, 235–242 (2011).

83. A. J. McCoy et al., Phaser crystallographic software. J Appl Crystallogr 40, 658–674 (2007).

84. P. Emsley, B. Lohkamp, W. G. Scott, K. Cowtan, Features and development of Coot. Acta Crystallogr D Biol Crystallogr 66, 486–501 (2010).

85. P. D. Adams et al., PHENIX: a comprehensive Python-based system for macromolecular structure solution. Acta Crystallogr D Biol Crystallogr 66, 213–221 (2010).

86. E. F. Pettersen et al., UCSF Chimera--a visualization system for exploratory research and analysis. J Comput Chem 25, 1605–1612 (2004).

87. R. A. Laskowski, M. B. Swindells, LigPlot+: multiple ligand-protein interaction diagrams for drug discovery. J Chem Inf Model 51, 2778–2786 (2011).

88. L. Holm, Benchmarking fold detection by DaliLite v.5. Bioinformatics 35, 5326–5327 (2019).

89. T. M. Ryan et al., An optimized SEC-SAXS system enabling high X-ray dose for rapid SAXS assessment with correlated UV measurements for biomolecular structure analysis. Applied Crystallography 51, 97–111 (2018).

90. N. Kirby et al., Improved radiation dose efficiency in solution SAXS using a sheath flow sample environment. Biological Crystallography 72, 1254–1266 (2016).

91. K. Manalastas-Cantos et al., ATSAS 3.0: expanded functionality and new tools for small-angle scattering data analysis. Applied Crystallography 54, 343–355 (2021).

92. A. Panjkovich, D. I. Svergun, CHROMIXS: automatic and interactive analysis of chromatography-coupled small-angle X-ray scattering data. Bioinformatics 34, 1944–1946 (2018).

93. J. B. Hopkins, BioXTAS RAW 2: new developments for a free open-source program for small-angle scattering data reduction and analysis. Applied Crystallography 57, 194–208 (2024).

94. D. Svergun, Determination of the regularization parameter in indirect-transform methods using perceptual criteria. Applied Crystallography 25, 495–503 (1992).

95. D. Franke, D. I. Svergun, DAMMIF, a program for rapid ab-initio shape determination in small-angle scattering. Applied Crystallography 42, 342–346 (2009).

96. V. V. Volkov, D. I. Svergun, Uniqueness of ab initio shape determination in small-angle scattering. Applied Crystallography 36, 860–864 (2003).

97. A. Panjkovich, D. I. Svergun, Deciphering conformational transitions of proteins by small angle X-ray scattering and normal mode analysis. Physical Chemistry Chemical Physics 18, 5707–5719 (2016).

98. Y. Kochnev, J. D. Durrant, FPocketWeb: protein pocket hunting in a web browser. J Cheminform 14, 58 (2022).

