## Supporting Information for "Unique structural and ligand binding properties of the *Staphylococcus aureus* serine hydrolase FphE"

**The PDF file includes:**

Figs. S1 to S13
Tables S1 to S13
Chemical synthesis and NMR spectra
References

**Other Supplementary Materials for this manuscript include the following:**

Supplementary Data 1 to 5

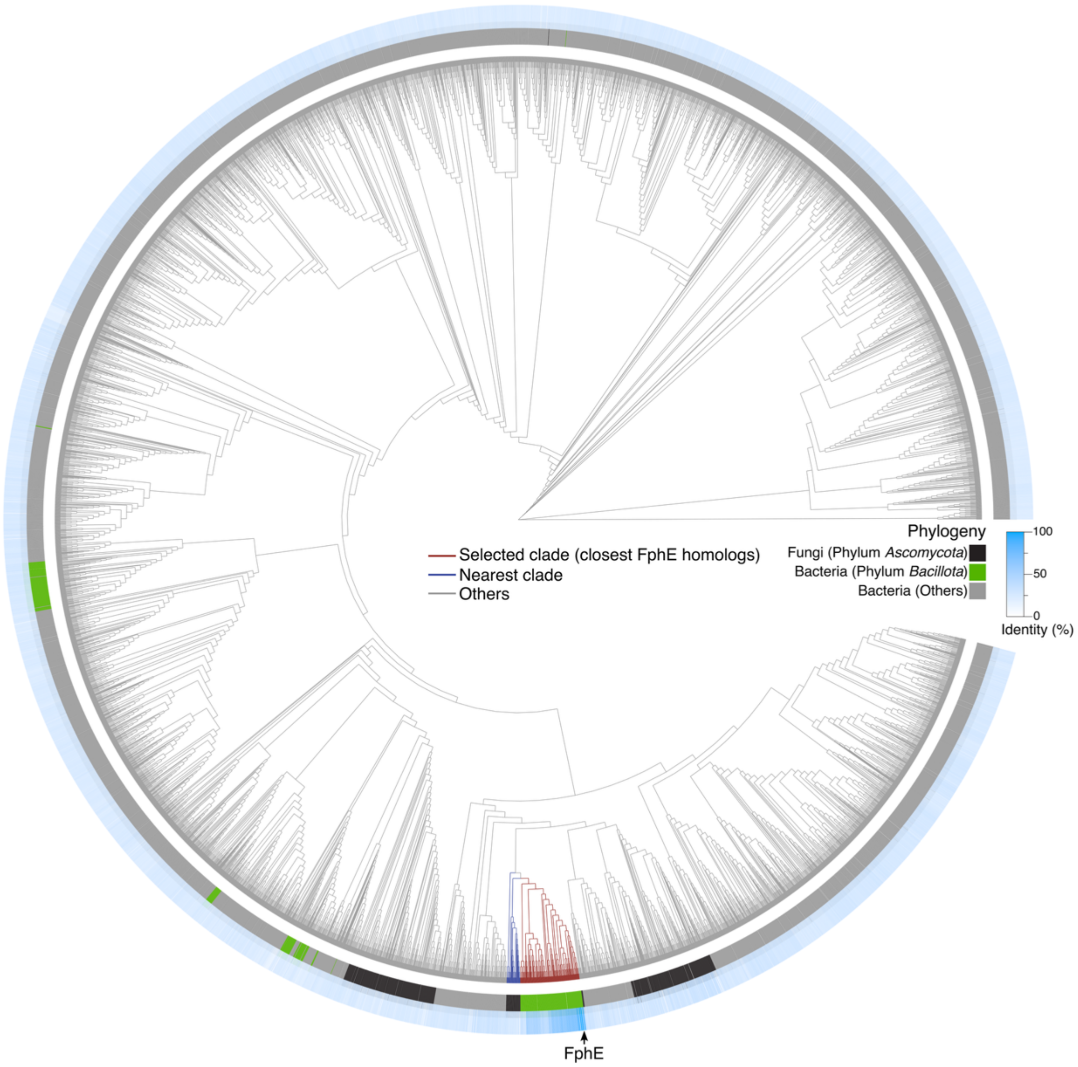


**Figure S1. A phylogenetic tree of 6,141 FphE-like proteins.** Rings are colored by phylogeny (inner) or sequence identity to FphE (outer).

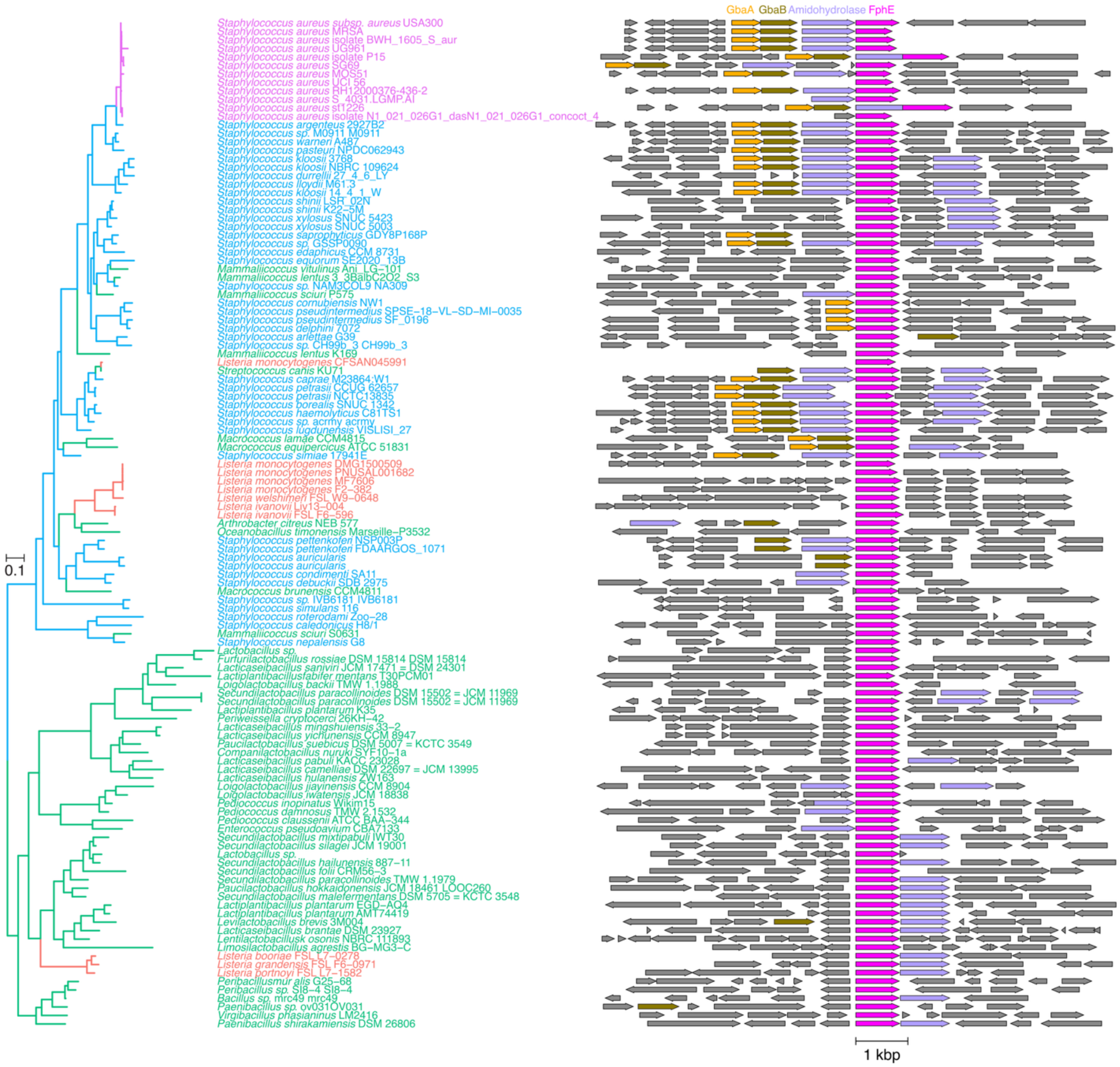


**Figure S2. A phylogenetic tree of 119 representative FphE homologs.** Representative species names and genomic loci are shown.

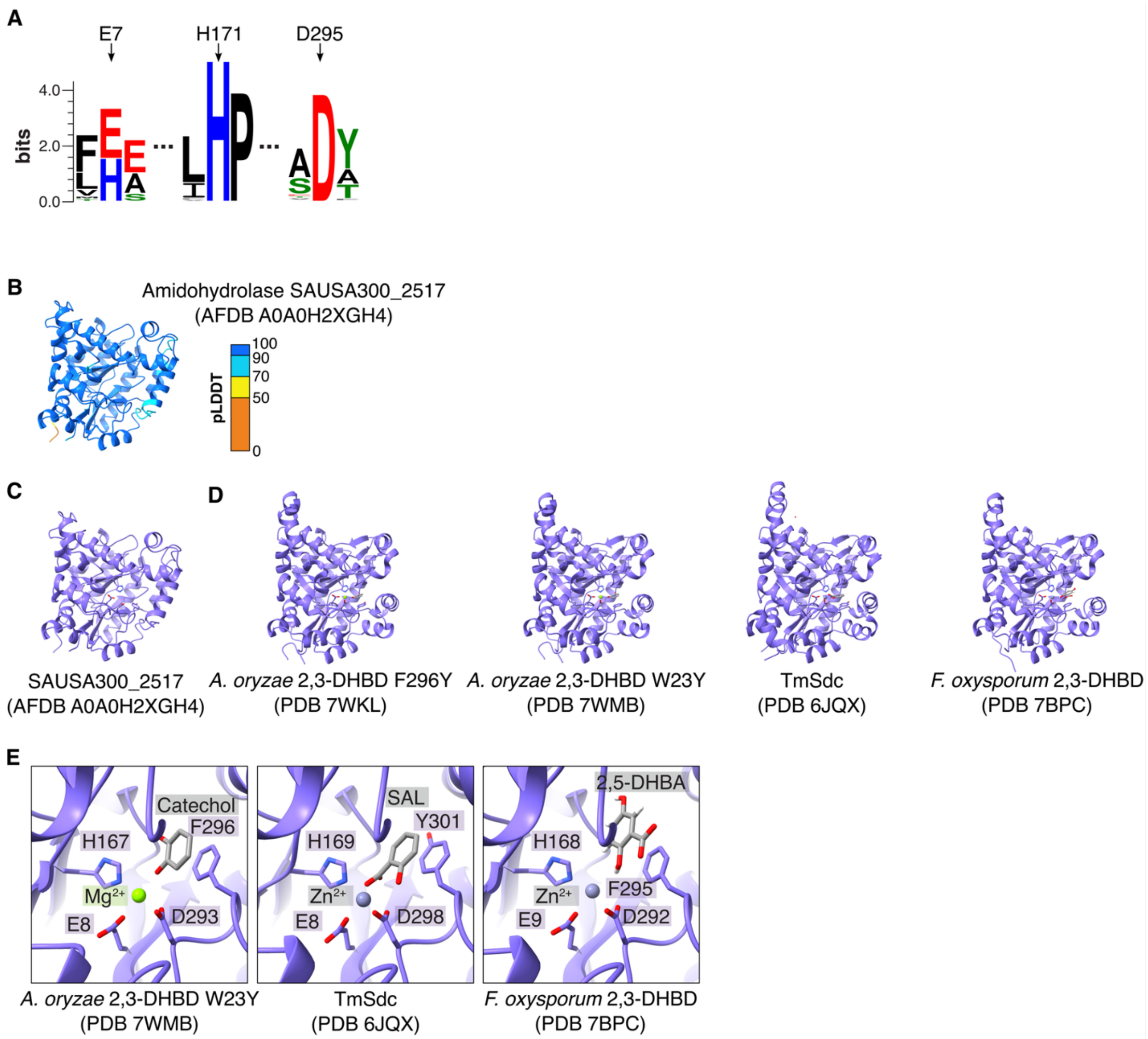


**Figure S3. Bioinformatic analysis of the putative amidohydrolase (SAUSA300_2517).** (**A**) Sequence logos for metal ion-binding catalytic residues of SAUSA300_2517 among 80 homologs in Fig. 1B. (**B–C**) Overall structure of AlphaFold2-predicted SAUSA300_2517 (AFDB A0A0H2XGH4) colored by either pLDDT score (A) or chain (B) (*1*). (**D–E**) Overall structures (D) and active sites (E) of *A. oryzae* 2,3-DHBD mutants (PDB 7WKL and 7WMB), *T. moniliiforme* salicylic acid decarboxylase (TmSdc; PDB 6JQX), and *F. oxysporum* 2,3-DHBD (PDB 7BPC) (*2-4*). 2,3-DHBD, 2,3-Dihydroxybenzoate decarboxylase; SAL, Salicylic acid; 2,5-DHBA, 2,5-Dihydroxybenzoic acid.

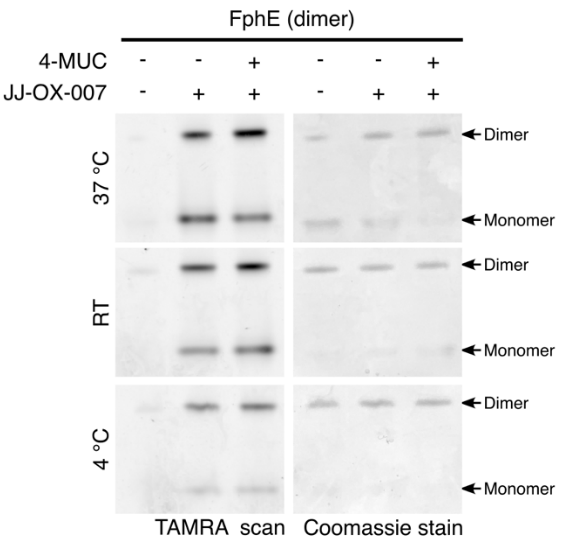


**Figure S4. Native-PAGE analysis of FphE dimer stability and probe labeling under variable temperatures.** FphE was incubated with or without 4-methylumbelliferyl caprylate (4-MUC) and/or the fluorescent at three different temperatures (37 °C, RT, and 4 °C), followed by native-PAGE analysis. RT, room temperature.

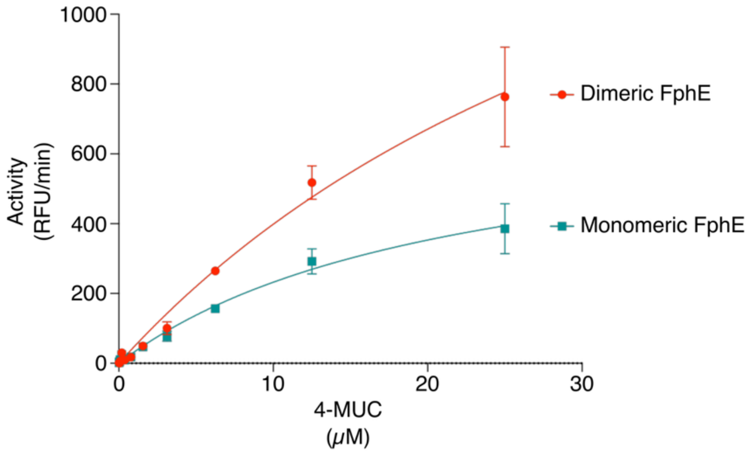


**Figure S5. Reaction rate for FphE monomer and dimer.** The progress curve of the FphE monomer and dimer protein (0.5 nM each) plotted in the presence of varied concentration range of 4-methylumbelliferone caprylate (4-MUC, 0–25μM). Data represent the three independent experiments and values are expressed in mean ± standard deviation.

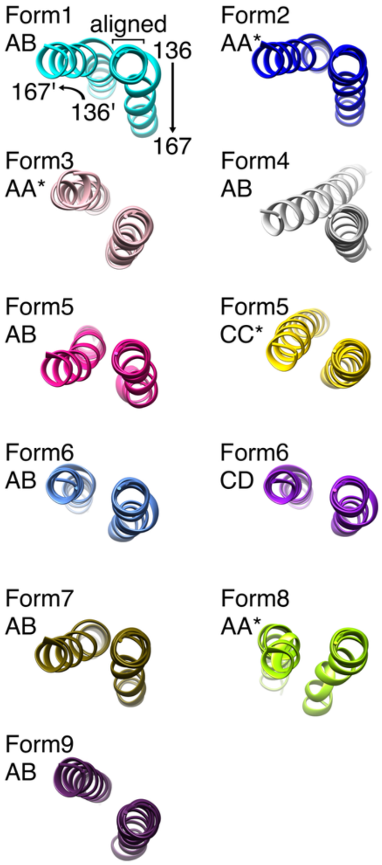


**Figure S6. Connecting helix 136–167 in different FphE crystal forms.** Connecting dimer helices 136–167 and dimer copy 136'–167' depicted in different ribbon colors. For Form 1 the aligned start of the first helix for all forms is indicated, as well as the ends of each helix. Form 1 chain A+B dimer cyan (PDB ID 8T87, dataset resolution 1.62 Å)(*5*); form 2 A+A* crystal symmetry dimer blue (8G49, 1.60 Å); form 3 A+A* pink (8G48, 1.95 Å); form 4 A+B grey (8UWM, 1.97 Å);(*6*) form 5 A+B magenta, C+C* yellow (9COM, 2.04 Å); form 6 A+B light blue, CD purple (9EBF, 2.50 Å); form 7 A+B ochre (8UIX, 2.39 Å);(*6*) form 8 A+A* green (9D87, 2.30 Å) and form 9 A+B dark purple (9EDJ, 2.60 Å).

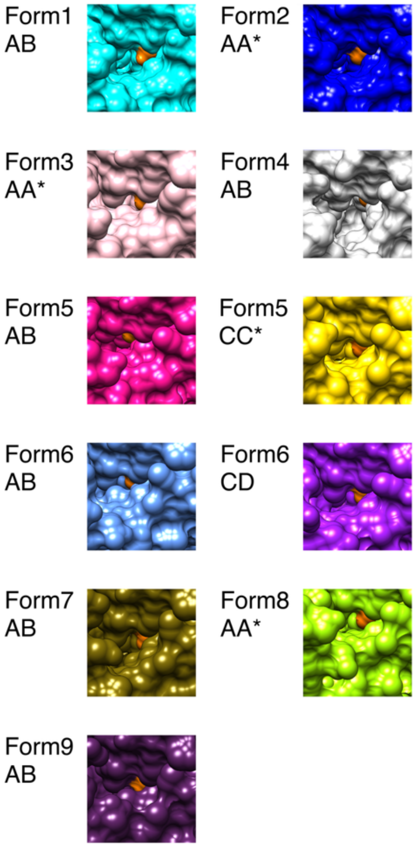


**Figure S7. Active site access in different FphE crystal forms.** Access to active site serine103 (orange surface depiction) for each crystal form in different surface colors. Form 1 chain A+B dimer cyan (PDB ID 8T87, dataset resolution 1.62 Å) (*5*); form 2 A+A* crystal symmetry dimer blue (8G49, 1.60 Å); form 3 A+A* pink (8G48, 1.95 Å); form 4 A+B grey (8UWM, 1.97 Å) (*6*); form 5 AB magenta, C+C* yellow (9COM, 2.04 Å); form 6 A+B light blue, C+D purple (9EBF, 2.50 Å); form 7 A+B ochre (8UIX, 2.39 Å) (*6*); form 8 A+A* green (9D87, 2.30 Å) and form 9 A+B dark purple (9EDJ, 2.60 Å).

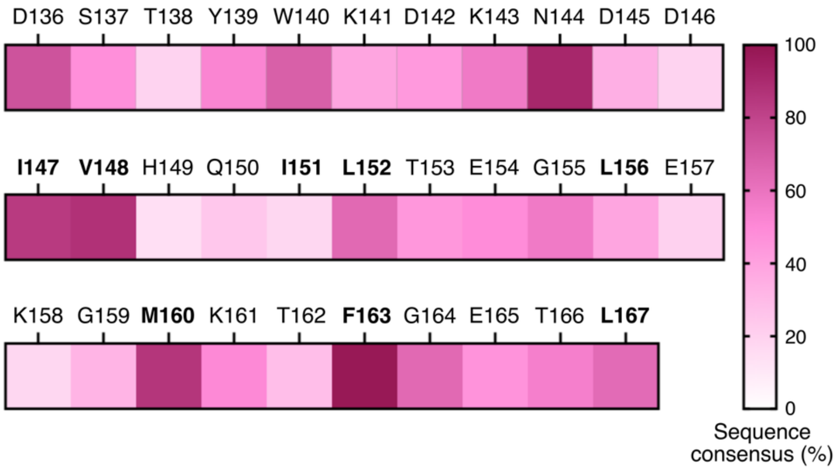


**Figure S8. Sequence conservation of residues 147–167 in FphE across 119 homologs.** The hydrophobic interfacing residues (Ile147, Val148, Ile151, Leu152, Leu156, Met160, Phe163 and Leu167) are annotated in bold.

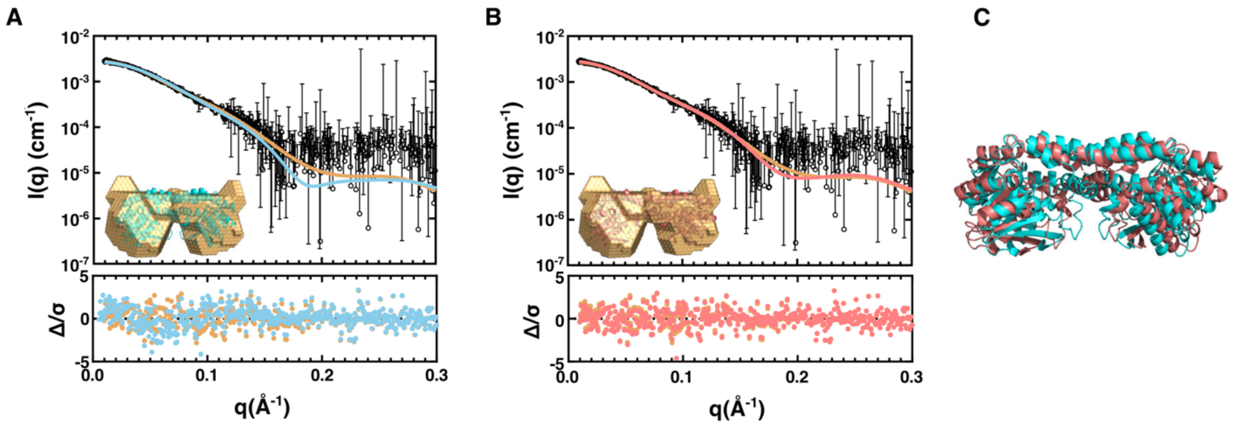


**Figure S9. Small-angle X-ray scattering (SAXS) of FphE.** (**A**) Top panel - experimental SAXS profile of FphE (black circles), with a fit model (beige line) used in DAMMIF to reconstruct an ab initio envelope (beige surface model) of FphE in solution. The theoretical scattering profile of the FphE crystal structure (PDB ID 8T87, blue line by Crysol) is overlaid. Insert ab initio envelope overlayed via ATSAS-CIFSUP with 8T87. Bottom panel residual error of the FphE crystal structure theoretical scattering curve (blue dots) and DAMMIF model (beige dots) versus q. (**B**) Experimental SAXS data are the same as in panel A compared to the SAXS-SREFLEX FphE model (red line, red cartoon model, and red dots) adjusted to better fit the data. (**C**) Overlay of the FphE crystal structure and SAXS-SREFLEX model.

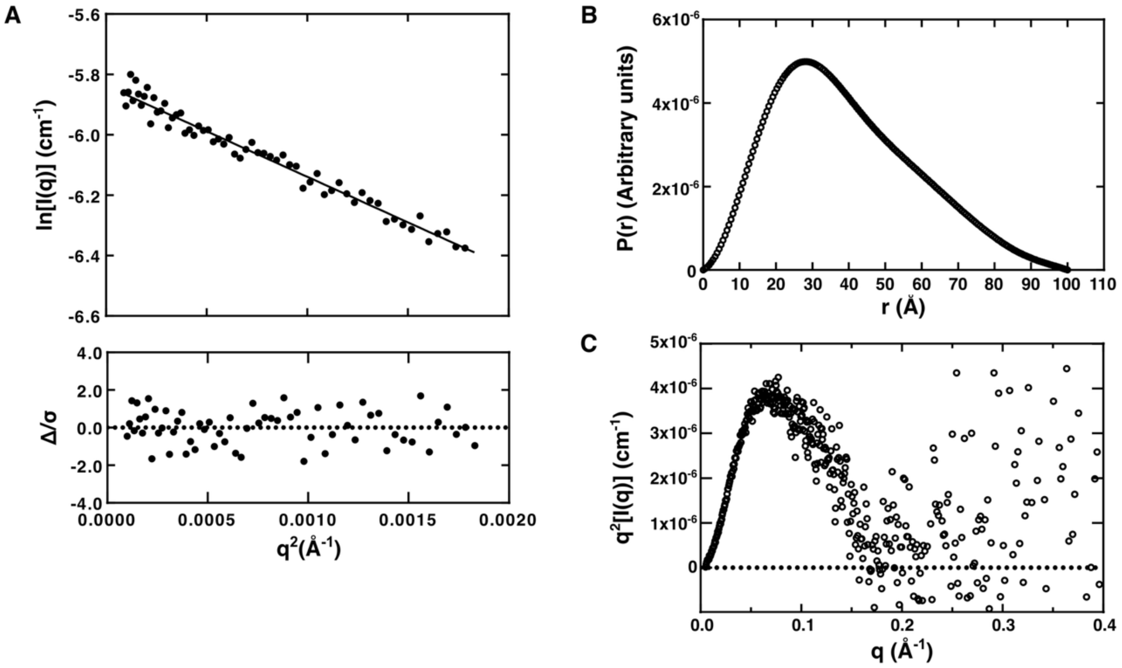


**Figure S10. Additional FphE SAXS graphs.** (**A**) Guinier plot (ln I(q) versus $q^2$) of low angle scattering, including the residuals ($\Delta/\sigma$) of the linear fit. (**B**) The distance distribution plot calculated using q to 0.3 $Å^{-1}$. (**C**) Kratky plot ($q^2I(q)$ versus q) of all data collected.

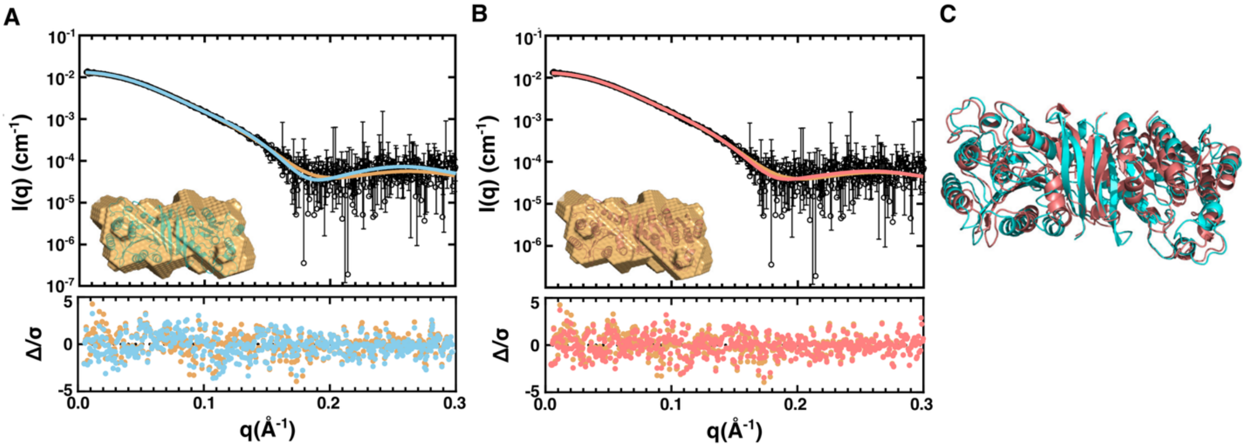


**Figure S11: Small-angle X-ray scattering (SAXS) of FphF.** (**A**) Top panel - experimental SAXS profile of FphF (black circles), with a fit model (beige line) used in DAMMIF to reconstruct an ab initio envelope (beige surface model) of FphF in solution. The theoretical scattering profile of the FphF crystal structure (PDB ID 6VH9 chain A and D, blue line by Crysol) is overlaid. Insert ab initio envelope overlayed via ATSAS-CIFSUP with 6VH9 chain A and D. Bottom panel residual error of the FphF crystal structure theoretical scattering curve (blue dots) and DAMMIF model (beige dots) versus q. (**B**) Experimental SAXS data are the same as in panel A compared to the SAXS-SREFLEX FphF model (red line, red cartoon model, and red dots) adjusted to better fit the data. (**C**) Overlay of the FphF crystal structure and SAXS-SREFLEX model.

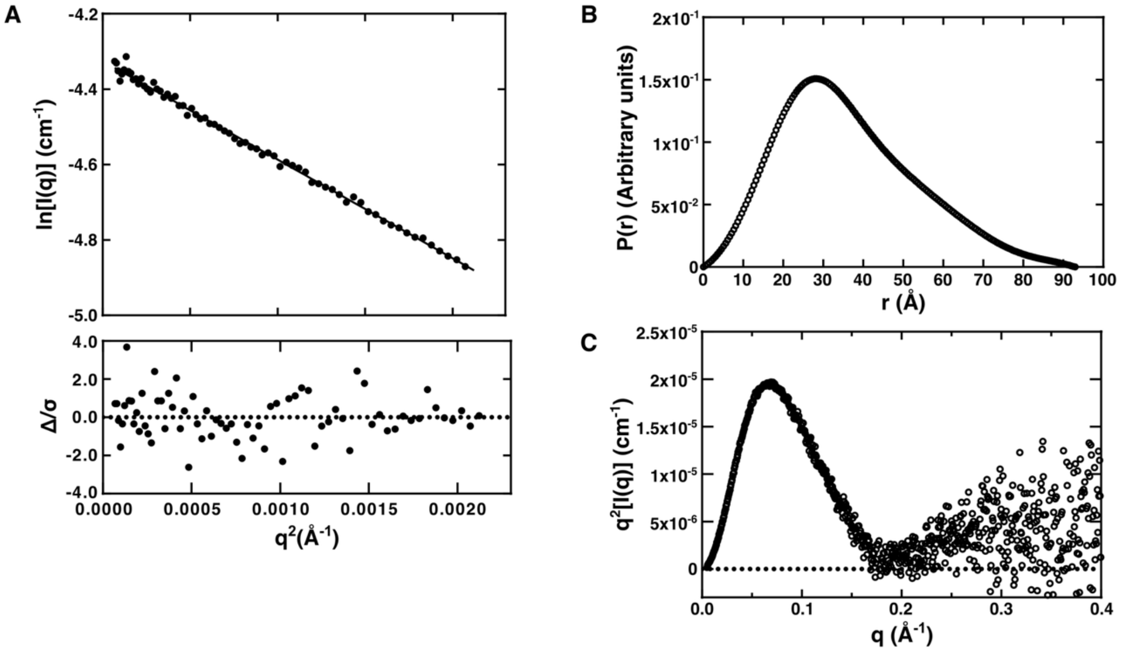


**Figure S12: Additional FphF SAXS graphs.** (**A**) Guinier plot (ln I(q) versus $q^2$) of low angle scattering, including the residuals (Δ/σ) of the linear fit. (**B**) The distance distribution plot calculated using q to 0.3 $Å^{-1}$. (**C**) Kratky plot ($q^2I(q)$ versus q) of all data collected.

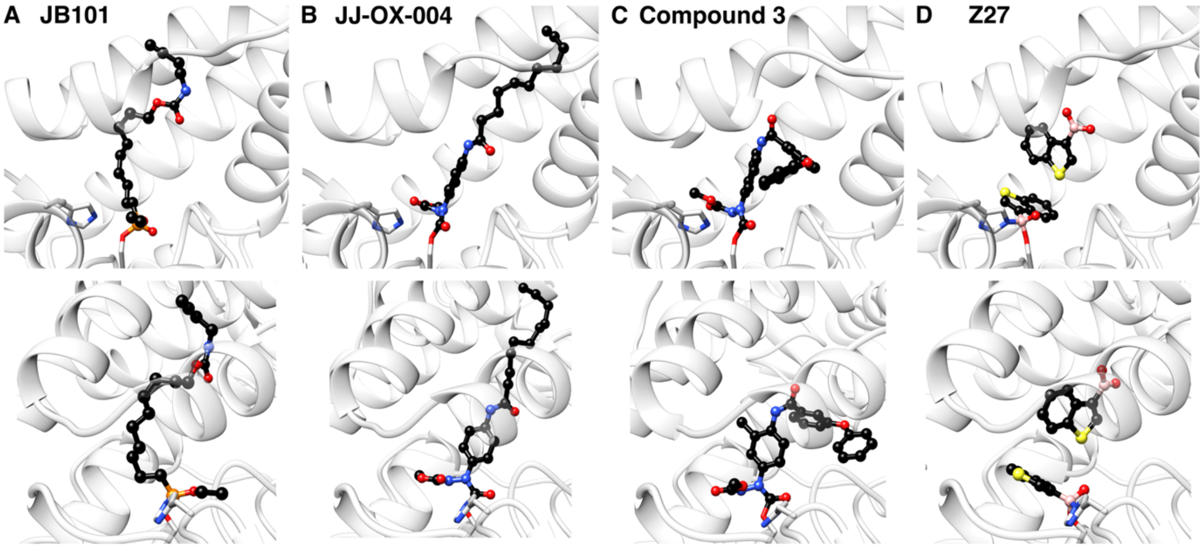


**Figure S13. Structural comparison of ligand binding to FphE.** Transparent grey ribbon representation of FphE, only side chains of active site triad residues Ser103 and His267 displayed. Covalent bound ligands in two orientations as balls and sticks with carbon atoms in black. (**A**) FP-JB101 (PDB ID 8SBQ, at 1.50 Å). (**B**) Oxadiazolone-JJ004 (8T88, 1.54 Å) (*5*). (**C**) Oxadiazolone-compound 3 (8G49, 1.60 Å). (**D**) Boronic acid-Z27 (8UGM, 1.65 Å) (*6*).

### Table S1. Representative genes for FphE homologs reported in previous papers.

See Supplementary Data 3 for the full list of FphE homologs identified in this study.

| UniProtKB Entry | Organism | Gene Names | Entry Name | RefSeq accessions | Ref |
|---|---|---|---|---|---|
| Q2FDS6 | Staphylococcus aureus (strain USA300) | SAUSA300_2518 | Y2518_STAA3 | WP_000448901.1 | (*5, 7-12*) |
| Q2FV39 | Staphylococcus aureus (strain NCTC 8325 / PS 47) | SAOUHSC_02900 | Y2900_STAA8 | WP_000448905.1 YP_501353.1; | (*12-17*) |
| Q2YWC8* | Staphylococcus aureus (strain bovine RF122 / ET3-1) | SAB2455 | Y2455_STAAB | WP_000448919.1 | (*18*) |
| Q5HCW9 | Staphylococcus aureus (strain COL) | SACOL2597 | Y2597_STAAC | WP_000448905.1 | (*19-25*) |
| Q7A3C4 | Staphylococcus aureus (strain N315) | SA2367 | Y2367_STAAN | WP_000448918.1 | (*18, 26-32*) |
| Q8NUP5 | Staphylococcus aureus (strain MW2) | MW2501 | Y2501_STAAW | WP_000448906.1 | (*33*) |
| Q99R57 | Staphylococcus aureus (strain Mu50 / ATCC 700699) | SAV2581 | Y2581_STAAM | WP_000448918.1 | (*26, 34*) |
| A0A0H3GIK6 | Listeria monocytogenes serotype 1/2a (strain 10403S) | LMRG_00440 | A0A0H3GIK6_LISM4 | WP_014600619.1 | (*35*) |
| A0A0H3KAP9 | Staphylococcus aureus (strain Newman) | NWMN_2480 | A0A0H3KAP9_STAAE | WP_000448905.1 | (*12, 27, 36*) |
| A0A3A2IXP9 | Staphylococcus aureus | DQV00_07015 G6Y24_05270 M1K003_1897 SAMEA70153168_01112 | A0A3A2IXP9_STAAU | WP_000448918.1 | (*26*) |
| A0A9P3ZH42 | Staphylococcus sp. 53017 | F1583_02530 | A0A9P3ZH42_9STAP | WP_000448918.1 | (*26*) |
| Q6GDM0 | Staphylococcus aureus (strain MRSA252) | SAR2661 | Y2661_STAAR | WP_000448898.1 | (*37*) |
| Q8Y8Z0 | Listeria monocytogenes serovar 1/2a (strain ATCC BAA-679 / EGD-e) | lmo0752 | Q8Y8Z0_LISMO | NP_464279.1 WP_003721869.1 | (*35, 38-41*) |

**Table S2. Model peptide substrates IQ1–IQ8 that were not cleaved by FphE.**

| | |
|---|---|
| IQ1 | Mca-His-Gln-Lys-Leu-Val-Phe-Phe-Ala-Lys(DNP)-NH2 |
| IQ2 | Mca-Glu-Val-Lys-Met-Asp-Ala-Glu-Phe-Lys(DNP)-NH2 |
| IQ3 | Mca-Ser-Glu-Val-Asn-Leu-Asp-Ala-Glu-Phe-Arg-Lys(DNP)-Arg-Arg-NH2 |
| IQ4 | Mca-Arg-Pro-Lys-Pro-Tyr-Ala-Nva-Trp-Met-Lys(DNP)-NH2 |
| IQ5 | Mca-Gly-Lys-Pro-Ile-Leu-Phe-Phe-Arg-Leu-Lys(DNP)-DArg-NH2 |
| IQ6 | Mca-Arg-Pro-Pro-Gly-Phe-Ser-Ala-Phe-Lys(DNP)-OH |
| IQ7 | Mca-Val-Asp-Gln-Met-Asp-Gly-Trp-Lys-(DNP)-NH2 |
| IQ8 | Abz-Gly-Ile-Val-Arg-Ala-Lys(DNP)-OH |

**Table S3. FphE monomer and dimer fractions quantified from native-PAGE (related to Fig. 2E and 2F).**

Values are reported as percentages calculated from integrated band intensities (Fig. 2E and 2F): Monomer (%) = 100 × $I_M/(I_M + I_D)$; Dimer (%) = 100 × $I_D/(I_M + I_D)$, where $I_M$ and $I_D$ are the integrated densities of the monomer and dimer bands, respectively. Data are shown as mean ± half-range (%), $n$ = 2. Because dimer (%) = 100 – monomer (%), the half-range of monomer and dimer fractions is identical. Half-range = (max − min)/2; N/A, not analyzed.

| Lane | Monomer fraction (Coomassie) | Monomer fraction (TAMRA) | Dimer fraction (Coomassie) | Dimer fraction (TAMRA) |
|---|---|---|---|---|
| 1 | 86.6 (1.2) | N/A | 13.4 (1.2) | N/A |
| 2 | 54.7 (1.2) | N/A | 45.3 (1.2) | N/A |
| 3 | 84.7 (4.4) | 80.0 (3.8) | 15.3 (4.4) | 20.0 (3.8) |
| 4 | 58.4 (4.7) | 56.5 (1.4) | 41.6 (4.7) | 42 (1.4) |

**Table S4. Crystal forms and crystallization conditions of fully processed FphE crystals**.

Highest diffracting crystal for each condition in Å.

| (Å) | Compound 1 | Compound 2 | Compound 3 |
|---|---|---|---|
| **Crystal form 1: P 1 21 2 or P 1 21 1; a, b, c (Å); α, β, γ (°) = ~47, 75, 75; 90, 91, 90; 2 chains** PDB: **8SBQ**, 8T87, 8T88, 8TFW, 8UGM, 9DRO | | | |
| 1.38 | 0.18 M Magnesium chloride | 0.1 M Tris pH 7.5/8.0 | 22.5 % w/v PEG 2000 MME |
| 1.65 | 0.2 M Magnesium chloride | 0.1 M Sodium acetate pH 5.5 | 10 % w/v PEG 8000<br>10 % w/v PEG 1000 |
| 1.65 | 0.18 M Calcium acetate | 0.1 M MES pH 6.5 | 22.5 % w/v PEG 2000 MME |
| 1.70 | 0.2 M Magnesium chloride | 0.1 M HEPES pH 7.4 | 25 % w/v PEG 2000 MME |
| 1.70 | 0.18 M Magnesium chloride | 0.1 M MES pH 6.5 | 22.5 % w/v PEG 2000 MME |
| 1.70 | 0.18 M Magnesium chloride | 0.1 M MIB pH 7.0 | 22.5 % w/v PEG 2000 MME |
| 1.83 | 0.16 M Magnesium chloride | | 25 % w/v PEG 2000 MME |
| 1.87 | 0.2 M Magnesium chloride | 0.1 M Tris pH 8.5 | 25 % w/v PEG 3350 |
| 1.98 | 0.2 M Magnesium formate | | 25 % w/v PEG 3350 |
| 2.00 | 0.18 M Magnesium chloride | 0.1 M Sodium acetate pH 5.5 | 22.5 % w/v PEG 2000 MME |
| 2.20 | 0.16 M Magnesium chloride | 0.18 M Tris pH 8.5 | 20 % w/v PEG 2000 MME |
| **Crystal form 2: P 32 2 1; a, b, c (Å); α, β, γ (°) = ~46, 46, 218; 90, 90, 120; 1 chain** PDB: **8G49** | | | |
| 1.60 | 0.16 M Potassium thiocyanate | 0.18 M Tris pH 8.5 | 20 % w/v PEG 2000 MME |
| **Crystal form 3: P 2 21 21; a, b, c (Å); α, β, γ (°) = ~42, 54, 122; 90, 90, 90; 1 chain** PDB: **8G48** | | | |
| 1.95 | 0.2 M Magnesium chloride | 0.1 M Tris pH 8.5 | 25 % w/v PEG 2000 MME |
| **Crystal form 4: P 21 21 21; a, b, c (Å); α, β, γ (°) = ~66, 78, 111; 90, 90, 90; 2 chains** PDB: 8UWM | | | |
| 1.97 | 0.18 M Calcium acetate | 0.1 M Tris pH 8.5 | 22.5 % w/v PEG 2000 MME |
| **Crystal form 5: P 2 21 21; a, b, c (Å); α, β, γ (°) = ~67, 87, 138; 90, 90, 90; 3 chains** PDB: **9COM** | | | |
| 2.04 | 0.2 M Potassium thiocyanate | 0.1 M Tris pH 8.5 | 25 % w/v PEG 2000 MME |
| 2.60 | 0.2 M Sodium acetate | 0.1 M Tris pH 8.5 | 25 % w/v PEG 2000 MME |
| **Crystal form 6: I 23; a, b, c (Å); α, β, γ (°) = ~212, 212, 212; 90, 90, 90; 4 chains** PDB: **9EBF** | | | |
| 2.50 | 0.18 M Potassium thiocyanate | 0.1 M Sodium acetate pH 5.5 | 22.5 % w/v PEG 2000 MME |
| **Crystal form 7: C 1 2 1; a, b, c (Å); α, β, γ (°) = ~47, 71, 165; 90, 96, 90; 2 chains** PDB: 8UIX | | | |
| 2.39 | 0.18 M Magnesium chloride | 0.1 M Tris pH 8.5 | 22.5 % w/v PEG 2000 MME |
| **Crystal form 8: P 23; a, b, c (Å); α, β, γ (°) = ~106, 106, 106; 90, 90, 90; 1 chain** PDB: **9D87** | | | |
| 2.30 | 0.18 M Potassium thiocyanate | 0.1 M Sodium acetate pH 5.5 | 22.5 % w/v PEG 2000 MME |
| **Crystal form 9: P 2 21 21; a, b, c (Å); α, β, γ (°) = ~52, 99, 102; 90, 90, 90; 2 chains** PDB: **9EDJ** | | | |
| 2.60 | 0.2 M Potassium thiocyanate | 0.1 M Sodium acetate pH 5.5 | 25 % w/v PEG 2000 MME |
| 3.10 | 0.18 M Magnesium chloride | 0.1 M MIB pH 7.0 | 22.5 % w/v PEG 2000 MME |
| 3.40 | 0.18 M Magnesium chloride | 0.1 M MMT pH 6.5 | 22.5 % w/v PEG 2000 MME |
| ~4 | 0.18 M Magnesium chloride | 0.1 M MES pH 6.5 | 22.5 % w/v PEG 2000 MME |
| **(Crystal form10 - twinning?: P 41 2 2; a, b, c (Å); α, β, γ (°) = ~70, 70, 99; 90, 90, 90; 1 chain?)** | | | |
| 2.42 | | 0.1 M MMT pH 6.0 | 25 % w/v PEG 1500 |
| 3.00 | 0.2 M Potassium bromide | 0.1 M Sodium acetate pH 5.5 | 25 % w/v PEG 2000 MME |
| **(Crystal form 11 - poor frames: P 2 2 21; a, b, c (Å); α, β, γ (°) = ~43, 77, 161; 90, 90, 90; 1 chain?)** | | | |
| 2.10 | 0.2 M Calcium acetate | 0.1 M MES pH 6.5 | 10 % w/v PEG 8000 |

| | | | |
|---|---|---|---|
| | | | 10 % w/v PEG 1000 |
| 2.65 | 0.2 M Calcium chloride | 0.1 M Tris pH 8.0 | 20 % w/v PEG 6000 |
| 2.68 | 0.2 M Magnesium chloride | 0.1 M Tris pH 8.5 | 25 % w/v PEG 3350 |
| **(<u>Crystal form 12 - tNCS:</u> P 42 21 2; a, b, c (Å); α, β, γ (°) = ~105, 105, 98; 90, 90, 90; 2 chains?)** | | | |
| 2.50 | 0.18 M Magnesium chloride | 0.1 M PCTP pH 7.0 | 22.5 % w/v PEG 2000 MME |
| 2.70 | 0.18 M Magnesium chloride | 0.1 M MES pH 6.5 | 22.5 % w/v PEG 2000 MME |
| 2.80 | 0.18 M Magnesium chloride | 0.1 M MIB pH 7.0 | 22.5 % w/v PEG 2000 MME |
| 2.93 | 0.18 M Magnesium chloride | | 22.5 % w/v PEG 2000 MME |
| 3.60 | 0.2 M Magnesium formate | pH 7.0 | 20 % w/v PEG 3350 |

PDB IDs bold and underlined reported in this work
PDB IDs 8T87, 8T88 reported previously – Jo et al (*5*).
PDB ID 9DRO reported previously – Wang et al (*42*).
PDB IDs 8TFW, 8UGM, 8UIX, 8UWM reported previously – Upadhyay et al (*6*).

**Table S5. Crystallization conditions of FphE not fully processed crystals.**

Highest diffracting crystal in Å. No dataset taken or processing failed early.

| (Å) | Compound 1 | Compound 2 | Compound 3 |
|---|---|---|---|
| ~2.0 | 0.2 M Sodium succinate | | 20 %w/v PEG 3350 |
| ~2.3 | 0.2 M Potassium nitrate | | 20 % w/v PEG 3350 |
| ~2.5 | 0.3 M Sodium acetate | 0.1 M Tris pH 8.5 | 10 % w/v PEG 8000<br>10 % w/v PEG 1000 |
| ~3 | 0.16 M Calcium acetate | 0.18 M MES pH 6.5 | 20 % w/v PEG 2000 MME |
| ~3 | 0.1 M Potassium thiocyanate | | 30 % w/v PEG 2000 MME |
| ~3 | 0.2 M Lithium chloride | | 20 % w/v PEG 3350 |
| ~3 | 0.2 M Potassium thiocyanate | | 20 % w/v PEG 3350 |
| ~3 | | 0.1 M PCB pH 6.0 | 25 % w/v PEG 1500 |
| ~3 | | 0.1 M MMT pH 6.0 | 25 % w/v PEG 1500 |
| ~3 | 0.3 M Sodium acetate | 0.1 M MES pH 6.5 | 25 % w/v PEG 2000 MME |
| ~3.2 | 0.15 M Potassium thiocyanate | 0.1 M Sodium acetate pH 5.5 | 18 % w/v PEG 5000 MME |
| ~3.2 | 0.2 M Ammonium acetate | 0.1 M Bis-Tris pH 5.5 | 25 % w/v PEG 3350 |
| ~3.4 | 0.8 M Sodium formate | 0.1 M TRIS pH 7.5 | 15 % w/v PEG 4000 |
| ~3.4 | 0.2 M Magnesium formate | | 25% w/v PEG 3350 |
| ~4 | 0.15 M Potassium thiocyanate | 0.1 M Sodium acetate pH 5.5 | 18 % w/v PEG 3350 |
| ~6 | 0.15 M Potassium thiocyanate | 0.1 M Sodium acetate pH 5.5 | 15 % w/v PEG 6000 |
| ~8 | 0.3 M Sodium acetate | 0.1 M Tris pH 8.5 | 25 % w/v PEG 2000 MME |
| ~10 | 0.2 M Sodium chloride | 0.1 M Tris pH 8.5 | 25 % w/v PEG 3350 |
| ~10 | 0.15 M Potassium bromide | | 30 % w/v PEG 2000 MME |

**Table S6. Crystallization conditions of FphE untested crystals.**

Diffraction potential not investigated.

| Compound 1 | Compound 2 | Compound 3 |
|---|---|---|
| 0.2 M Calcium acetate | | 25 % w/v PEG 2000 MME |
| 0.2 M Calcium acetate | 0.1M HEPES pH 7.4 | 25 % w/v PEG 2000 MME |
| 0.2 M Calcium acetate | 0.1 M Tris pH 8.0 | 25 % w/v PEG 2000 MME |
| 0.8 M Sodium formate | 0.1 M TRIS pH 7.5 or 8.5 | 25 % w/v PEG 2000 MME |
| 0.2 M Potassium thiocyanate | | 25 % w/v PEG 2000 MME |
| 0.2 M Potassium thiocyanate | 0.1 M MES pH 6.5 | 25 % w/v PEG 2000 MME |
| 0.2 M Potassium thiocyanate | 0.1M HEPES pH 7.4 | 25 % w/v PEG 2000 MME |
| 0.2 M Potassium thiocyanate | 0.1 M Tris pH 7.5 or 8.0 | 25 % w/v PEG 2000 MME |
| 0.2 M Potassium bromide | 0.1 M MES pH 6.5 | 25 % w/v PEG 2000 MME |
| 0.2 M Potassium bromide | 0.1 M Tris pH 7.5 or 8.5 | 25 % w/v PEG 2000 MME |
| 0.3 M Sodium acetate | | 25 % w/v PEG 2000 MME |
| 0.3 M Sodium acetate | 0.1 M Sodium acetate pH 5.5 | 25 % w/v PEG 2000 MME |
| 0.3 M Sodium acetate | 0.1M HEPES pH 7.4 | 25 % w/v PEG 2000 MME |
| 0.3 M Sodium acetate | 0.1 M Tris pH 7.5 | 25 % w/v PEG 2000 MME |
| 0.2 M Calcium acetate | 0.1 M MES pH 6.5 | 15 % w/v PEG 4000 |
| 0.8 M Sodium formate | 0.1 M Tris pH 7.5 or 8.5 | 15 % w/v PEG 4000 |
| 0.005 M Cadmium chloride | 0.1 M Tris pH 8.5 | 20 % w/v PEG 4000 |
| 0.3 M Sodium acetate | 0.1 M MES pH 6.5 | 10 % w/v PEG 8000<br>10 % w/v PEG 1000 |
| 0.3 M Sodium acetate | 0.1 M TRIS pH 7.5 | 10 % w/v PEG 8000<br>10 % w/v PEG 1000 |
| 0.8 M Sodium formate | 0.1 M TRIS pH 8.5 | 10 % w/v PEG 8000<br>10 % w/v PEG 1000 |
| 0.2 M Potassium bromide | 0.1 M TRIS pH 7.5 | 10 % w/v PEG 8000<br>10 % w/v PEG 1000 |
| 0.2 M Magnesium chloride | 0.1 M TRIS pH 8.5 | 10 % w/v PEG 8000<br>10 % w/v PEG 1000 |
| 0.2 M Calcium acetate | 0.1 M Sodium acetate pH 5.5 | 10 % w/v PEG 8000<br>10 % w/v PEG 1000 |
| 0.2 M Potassium bromide | 0.1 M Sodium acetate pH 5.5 | 10 % w/v PEG 8000<br>10 % w/v PEG 1000 |
| 0.2 M Calcium acetate | 0.1 M Sodium acetate pH 5.5 | 8 % w/v PEG 20000<br>8 % v/v PEG 550 MME |
| 0.1 M Sodium succinate | | 15 % w/v PEG 3350 |
| 0.15 M CsCl | | 15 % w/v PEG 3350 |
| 0.01 M Magnesium chloride<br>0.005 M Nickel(II) chloride | 0.1 M HEPES sodium pH 7.0 | 15% w/v PEG 3350 |
| 0.15 M Potassium thiocyanate | 0.1 M MES pH 6.5 | 18 % w/v PEG 3350 |
| 0.15 M Potassium thiocyanate | 0.1 M Tris pH 7.5 | 18 % w/v PEG 3350 |
| 0.15 M Potassium thiocyanate | 0.1 M Sodium acetate pH 5.5 | 18 % w/v PEG 3350 |
| 0.2 M Sodium formate | | 20 % w/v PEG 3350 |
| 0.2 M Ammonium formate | | 20 % w/v PEG 3350 |
| 0.2 M Potassium formate | | 20 % w/v PEG 3350 |
| 0.2 M Potassium nitrate | | 20 % w/v PEG 3350 |
| 0.2 M Potassium fluoride | | 20 % w/v PEG 3350 |
| 0.2 M Ammonium fluoride | | 20 % w/v PEG 3350 |
| 0.2 M Potassium acetate | | 20 % w/v PEG 3350 |

| | | |
|---|---|---|
| 2 % v/v Tacsimate | 0.1 M HEPES pH 7.5 | 20 % w/v PEG 3350 |
| 0.01 M Magnesium chloride | 0.005 M NiCl2 | 20 % w/v PEG 3350 |
| 0.2 M Sodium chloride | 0.1 M Bis-Tris pH 5.5 | 25 % w/v PEG 3350 |
| 0.2 M Ammonium acetate | 0.1 M Bis-Tris pH 5.5 | 25 % w/v PEG 3350 |
| 0.2 M Ammonium acetate | 0.1 M Bis-Tris pH 5.5 | 25 % w/v PEG 3350 |
| 0.2 M Sodium chloride | 0.1 M Tris pH 8.5 | 25 % w/v PEG 3350 |
| 0.2 M Magnesium chloride | 0.1 M Tris pH 8.5 | 25 % w/v PEG 3350 |
| 0.2 M Magnesium formate | | 25 % w/v PEG 3350 |
| 0.15 M Potassium thiocyanate | 0.1 M Sodium acetate pH 5.5 | 20 % w/v PEG 1500 |
| 0.15 M Potassium thiocyanate | 0.1 M Tris pH 7.5 | 20 % w/v PEG 1500 |
| 0.15 M Potassium thiocyanate | 0.1 M Tris pH 8.5 | 20 % w/v PEG 1500 |
| | 0.1 M SPG pH 5.0 | 25 % w/v PEG 1500 |
| | 0.1 M MIB pH 6.0 | 25 % w/v PEG 1500 |
| | 0.1 M MIB pH 7.0 | 25 % w/v PEG 1500 |
| | 0.1 M PCB pH 5.0 | 25 % w/v PEG 1500 |
| | 0.1 M MMT pH 7.0 | 25 % w/v PEG 1500 |
| 0.15 M Potassium thiocyanate | 0.1 M Sodium acetate pH 5.5 | 18 % w/v PEG 5000 MME |
| 0.15 M Potassium thiocyanate | 0.1 M Tris pH 8.5 | 18 % w/v PEG 5000 MME |
| 0.15 M Potassium thiocyanate | 0.1 M Tris pH 7.5 | 18 % w/v PEG 5000 MME |
| 0.15 M Potassium thiocyanate | 0.1 M MES pH 6.5 | 18 % w/v PEG 5000 MME |
| 0.02 M Magnesium chloride | 0.1 M HEPES pH 7.5 | 22 % w/v PAA 5100 |
| | 0.1 M Sodium acetate pH 5.5 | 1.5 M Ammonium sulfate |
| | 0.1 M Tris pH 7.5 | 1.5 M Ammonium sulfate |
| | 0.1 M Sodium acetate pH 5.5 | 30 % v/v Jeffamine M-600 |
| | 0.1 M Tris pH 8.5 | 0.7 M Sodium citrate |
| | 0.1 M Bis-Tris prop pH 7.0 | 1.2 M Sodium citrate |
| | 0.1 M Tris pH 8.5 | 1.2 M Sodium citrate |

### Table S7. FphE unbound and ligand data collection and processing.

Values for the outer shell are given in parentheses.

| | JB101 bound in crystal form 1 | Compound-3 bound in crystal form 2 | Unbound crystal form 3 | Unbound crystal form 5 | Q41 bound in crystal form 6 | Unbound crystal form 8 | Unbound crystal form 9 |
|---|---|---|---|---|---|---|---|
| PDB ID | 8SBQ | 8G49 | 8G48 | 9COM | 9EBF | 9D87 | 9EDJ |
| Diffraction source | Australian synchrotron MX2 | Australian synchrotron MX2 | Australian synchrotron MX2 | Australian synchrotron MX2 | Australian synchrotron MX2 | Australian synchrotron MX1 | Australian synchrotron MX2 |
| Wavelength (Å) | 0.954 | 0.954 | 0.954 | 0.954 | 0.954 | 0.954 | 0.954 |
| Detector | DECTRIS EIGER X 16M | DECTRIS EIGER X 16M | DECTRIS EIGER X 16M | DECTRIS EIGER X 16M | DECTRIS EIGER X 16M | DECTRIS EIGER2 9M | DECTRIS EIGER X 16M |
| Space group | $P\ 1\ 2_1\ 1$ | $P\ 3_2\ 2\ 1$ | $P\ 2\ 2_1\ 2_1$ | $P\ 2\ 2_1\ 2_1$ | I 2 3 | P 2 3 | $P\ 2\ 2_1\ 2_1$ |
| a, b, c (Å) | 46.9 74.6 73.9 | 46.4 46.4 218.4 | 41.9 53.8 121.6 | 67.4 86.8 137.8 | 212.3 212.3 212.3 | 105.7 105.7 105.7 | 52.3 98.8 102.4 |
| α, β, γ (°) | 90.0 91.5 90.0 | 90.0 90.0 120.0 | 90.0 90.0 90.0 | 90.0 90.0 90.0 | 90.0 90.0 90.0 | 90.0 90.0 90.0 | 90.0 90.0 90.0 |
| Resolution range (Å) | 46.70 – 1.49 (1.52 – 1.49) | 43.67 – 1.60 (1.63 – 1.60) | 41.94 – 1.95 (2.00 – 1.95) | 48.17 – 2.04 (2.10 – 2.04) | 47.47 – 2.50 (2.57 – 2.50) | 47.27 – 2.30 (2.38 – 2.30) | 46.59 – 2.60 (2.72 – 2.60) |
| Total No. of reflections | 1076479 (30839) | 505683 (12794) | 126180 (8309) | 250519 (18545) | 1710359 (135026) | 539288 (50595) | 104515 (12542) |
| No. of unique reflections | 81242 (3577) | 37428 (1700) | 20746 (1384) | 51434 (3773) | 54846 (4421) | 17790 (1680) | 16871 (1962) |
| Completeness (%) | 99.4 (88.6) | 99.7 (95.6) | 99.5 (95.2) | 99.2 (95.2) | 99.9 (98.4) | 99.8 (98.4) | 99.6 (97.3) |
| Redundancy | 13.3 (8.6) | 13.5 (7.5) | 6.1 (6.0) | 4.9 (4.9) | 31.2 (30.5) | 30.3 (30.1) | 6.2 (6.4) |
| $\langle I/\sigma(I) \rangle$ | 8.0 (1.2) | 15.0 (1.4) | 9.7 (2.1) | 11.9 (1.2) | 23.7 (2.1) | 32.8 (1.6) | 9.8 (2.4) |
| $CC_{1/2}$ | 0.997 (0.643) | 0.999 (0.581) | 0.997 (0.717) | 0.999 (0.522) | 1.000 (0.692) | 1.000 (0.654) | 0.998 (0.860) |
| $R_{merge.}$ | 0.143 (1.116) | 0.079 (1.208) | 0.093 (0.758) | 0.062 (1.338) | 0.121 (2.141) | 0.079 (2.712) | 0.104 (0.762) |
| $R_{p.i.m.}$ | 0.059 (0.558) | 0.031 (0.675) | 0.060 (0.492) | 0.046 (0.984) | 0.031 (0.552) | 0.020 (0.700) | 0.067 (0.477) |

### Table S8. FphE unbound and ligand structure solution and refinement.

Values for the outer shell are given in parentheses.

| | JB101 bound in crystal form 1 | Compound-3 bound in crystal form 2 | Unbound crystal form 3 | Unbound crystal form 5 | Q41 bound in crystal form 6 | Unbound crystal form 8 | Unbound crystal form 9 |
|---|---|---|---|---|---|---|---|
| PDB ID | 8SBQ | 8G49 | 8G48 | 9COM | 9EBF | 9D87 | 9EDJ |
| Resolution range (Å) | 40.09 – 1.50 (1.52 – 1.50) | 39.55 – 1.60 (1.64 – 1.60) | 41.94 – 1.95 (2.05 – 1.95) | 43.40 – 2.04 (2.09 – 2.04) | 47.47 – 2.50 (2.54 – 2.50) | 37.37 – 2.30 (2.44 – 2.30) | 46.59 – 2.60 (2.76 – 2.60) |
| Final R-work | 0.166 (0.311) | 0.175 (0.321) | 0.182 (0.236) | 0.197 (0.331) | 0.183 (0.308) | 0.218 (0.331) | 0.188 (0.248) |
| Final R-free | 0.189 (0.329) | 0.198 (0.341) | 0.227 (0.309) | 0.248 (0.313) | 0.222 (0.326) | 0.237 (0.357) | 0.268 (0.248) |
| Protein residues | 554 | 276 | 277 | 833 | 1112 | 277 | 554 |
| Ligands | 2 (JB101) | 1 (Cmp3) | 0 | 0 | 4 (Q41) | 0 | 0 |
| Water | 534 | 183 | 137 | 206 | 76 | 12 | 24 |
| Metals | 3 (Mg) | 0 | 0 | 1 (K) | 0 | 0 | 0 |
| R.m.s. deviations | | | | | | | |
| Bonds (Å) | 0.006 | 0.011 | 0.007 | 0.008 | 0.009 | 0.008 | 0.008 |
| Angles (°) | 0.914 | 1.228 | 0.947 | 0.912 | 1.008 | 0.999 | 1.002 |
| Average *B* factors (Å$^2$) | 28.6 | 35.8 | 34.6 | 57.7 | 72.5 | 75.6 | 62.7 |
| Ligands | 37.6 | 32.5 | - | - | 62.99 | - | - |
| Water | 36.5 | 39.2 | 40.0 | 48.8 | 61.4 | 66.8 | 54.2 |
| Metals | 26.1 | - | - | 56.6 | - | - | - |
| Ramachandran plot | | | | | | | |
| Most favored (%) | 98.7 | 98.9 | 98.6 | 97.9 | 97.2 | 95.3 | 96.9 |
| Outlier (%) | 0 | 0 | 0 | 0 | 0 | 0 | 0 |

### Table S9. FphE dimer connecting helix distances.

Distances of Cα to Cα atom of both ends of the helix (137-167), as well as the middle residue (155) indicated for each crystal form.

| Form | PDB | Dimer | 137-167 [Å] | 167-137 [Å] | 155-155 [Å] |
|---|---|---|---|---|---|
| 1 | 8T87 | A+B | 13.3 | 13.7 | 6.0 |
| 2 | 8G49 | A+A* | 14.0 | 14.0 | 5.2 |
| 3 | 8G48 | A+A* | 13.4 | 13.4 | 8.2 |
| 4 | 8UWM | A+B | 19.2 | 19.0 | 4.5 |
| 5 | 9COM | A+B | 14.8 | 13.4 | 6.6 |
| 5 | 9COM | C+C* | 13.5 | 13.5 | 5.9 |
| 6 | 9EBF | A+B | 16.6 | 16.5 | 10.8 |
| 6 | 9EBF | C+D | 16.7 | 16.7 | 10.8 |
| 7 | 8UIX | A+B | 13.6 | 13.7 | 4.2 |
| 8 | 9D87 | A+A* | 13.1 | 13.1 | 6.9 |
| 9 | 9EDJ | A+B | 14.4 | 14.1 | 7.0 |

### Table S10. Parameters for Small-Angle X-ray Scattering of FphE.

| Data collection parameters | FphE SASBDB ID SASDX64 | FphF SASBDB ID SASDX74 |
|---|---|---|
| Beamline | Australian Synchrotron BioSAXS beamline | Australian Synchrotron BioSAXS beamline |
| Wavelength | 1.00 Å | 1.00 Å |
| Detector | Pilatus3X-2M | Pilatus3X-2M |
| Camera Length | 2.5m | 2.5m |
| SEC Column | Superdex Increase 75 5/150 GL | Superdex Increase 75 5/150 GL |
| q range | 0.00502665-0. 551690 | 0.00502665-0.551690 |
| sample capillary flow rate | 0.4 mL/min | 0.4 mL/min |
| Exposure time | 1 second | 1 second |
| Number of Frames used | 20 | 15 |
| Sample concentration injected | 1.0 mg/ml | 6.5 mg/ml |
| Sample volume injected | 50 µl | 50 µl |
| Temperature | Room temperature ~21 °C | Room temperature ~21 °C |
| Structural parameters | | |
| q range for P(r) ($Å^{-1}$) | 0.01-0.3 | 0.005-0.3 |
| I (0) from P(r) ($cm^{-1}$) | 0.00292 ± 0.0000208 | 0.0132±0.0000215 |
| Rg (Å) from P(r) (Å) | 30.7400±0.2441 | 28.28±0.0601 |
| q range for Guinier ($Å^{-1}$) | 0.00995-0.04278 | 0.00831-0.04607 |
| I(0) from Guinier ($cm^{-1}$) | 0.00292±0.0000217 | 0.0132±0.0000232 |
| Rg (Å) from Guinier (Å) | 30.0805± 0.3618 | 28.0158±0.0809 |
| Dmax (Å) | 100 | 93 |
| Porod volume estimate ($Å^3$) | 79800 | 76200 |
| MW from sequence* | 31,242 Da | 29,307 Da |
| MW from Bayesian Inference | 62,400 Da | 55,600 Da |
| MW Probability from Bayesian Inference | 0.28 | 0.269 |
| Credibility Interval from Bayesian Inference | 57,500-69,700 Da | 51,500-58,800 Da |
| Credibility Interval Probability from Bayesian Inference | 0.916 | 0.972 |
| Software | | |
| Data Processing | pyFAI | pyFAI |
| Buffer Subtraction | chromixs, ATSAS 3.2.2 | chromixs, ATSAS 3.2.2 |
| Data Analysis | RAW/Primus/GraphPad Prism | RAW/Primus/GraphPad Prism |

*Full length sequence of FphE (SAUSA300_2518) and FphF (SAUSA300_2564) with N-terminal GPG from expression tag after 3C cleavage.

**Table S11. Protein structures similar to predicted FphE monomer structure.**

Top5 Dali (*43*) hits using *S. aureus* USA300 LAC Q2FDS6 Uniprot (*44*) predicted Alphafold 2 (*45*) AF-Q2FDS6-F1-model_v4 with representative PDB IDs from a PDB25 search.

| # | Representative PDB entry ID | Z | rmsd | lali | nres | % id | Description |
|---|---|---|---|---|---|---|---|
| **1** | 5XMW | 24.8 | 3.1 | 247 | 270 | 18 | Zearalenone lactonase from *Clonostachys rosea* |
| **2** | 3R0V | 23.7 | 3.4 | 245 | 257 | 22 | Alpha/beta hydrolase from *Sphaerobacter thermophilus* |
| **3** | 4Q3L | 22.5 | 3.1 | 244 | 280 | 16 | Alpha/beta hydrolase from deep-sea metagenome |
| **4** | 4QES | 22.5 | 3.2 | 243 | 441 | 17 | Alpha/beta hydrolase from *Kitasatospora aureofacien* |
| **5** | 4L0C | 22.2 | 3.0 | 221 | 255 | 19 | Alpha/beta hydrolase from *Pseudomonas putida* S16 |

**Table S12. Protein structures similar to FphE dimer chain A in crystal from 1.**

Top5 Dali (*43*) hits using chain A of FphE structure 8T87 with representative PDB IDs from a PDB25 search.

| # | Representative PDB entry ID | Z | rmsd | lali | nres | % id | Description |
|---|---|---|---|---|---|---|---|
| **1** | 7D78 | 16.3 | 3.2 | 143 | 297 | 20 | Thioesterase from *Beauveria bassiana* |
| **2** | 7C4D | 16.0 | 3.9 | 137 | 261 | 29 | Alpha/beta hydrolase from uncultured bacterium |
| **3** | 4L0C | 15.9 | 3.9 | 136 | 255 | 23 | Alpha/beta hydrolase from *Pseudomonas putida* S16 |
| **4** | 6T6H | 15.8 | 3.6 | 140 | 253 | 16 | Bottromycin epimerase from *Streptomyces sp.* BC16019 |
| **5** | 7AL6 | 15.7 | 3.5 | 141 | 285 | 24 | Alpha/beta hydrolase from *Pseudomonas aeruginosa* PAO1 |

**Table S13.** ***Staphylococcus aureus*** **predicted AlphaFoldDB proteome (STAA8) similar to FphE dimer chain A in crystal from 1.**

Top5 Dali (*43*) hits using chain A of FphE structure 8T87 searched against the predicted *Staphylococcus aureus* proteome via AlphaFoldDB (*45*).

| # | AlphaFoldDB ID | Z | rmsd | lali | nres | % id | Description |
|---|---|---|---|---|---|---|---|
| **1** | AF-Q2FV39-F1 | 19.3 | 1 | 155 | 276 | 100 | FphE |
| **2** | AF-Q2G0F0-F1 | 13.2 | 3.3 | 139 | 262 | 22 | Uncharacterized ABhydrolase_1 |
| **3** | AF-Q2G0F3-F1 | 12.8 | 4 | 138 | 266 | 23 | Uncharacterized ABhydrolase_1 |
| **4** | AF-Q2FV24-F1 | 11.3 | 9.6 | 132 | 560 | 17 | Uncharacterized Xaa-Pro dipeptidyl-peptidase |
| **5** | AF-Q2G2V6-F1 | 11.2 | 2.7 | 127 | 248 | 14 | FphG |

### Chemical Synthesis and NMR spectra

#### General methods

Unless noted otherwise, reagents and solvents were purchased from commercial suppliers and were used without further purification. All reactions were performed under argon atmosphere and stirring unless otherwise noted. Reaction flasks were dried overnight at 100 °C in an oven. Flash column chromatography was carried out using SiliaFlash® P60 (230–400 mesh, SiliCycle) with the indicated solvents of reagent-grade. Thin-layer chromatography (TLC) was performed using pre-coated 0.25 mm silica gel plates (Merck). $^{1}$H and $^{13}$C NMR spectra were recorded on Varian 400 spectrometer as solutions in the indicated solvents at room temperature (rt). Chemical shifts were quoted in parts per million (ppm, δ) downfield from tetramethylsilane (TMS) and were referenced to the deuterated solvent. $^{1}$H NMR data were reported in the order of chemical shift, multiplicity (s, singlet; d, doublet; t, triplet; q, quartet; quint, quintet; m, multiplet and/or multiple resonance), number of protons, and coupling constant in hertz (Hz). The following abbreviation for solvents are used: *N*,*N*-dimethylformamide (DMF), dimethyl sulfoxide (DMSO).

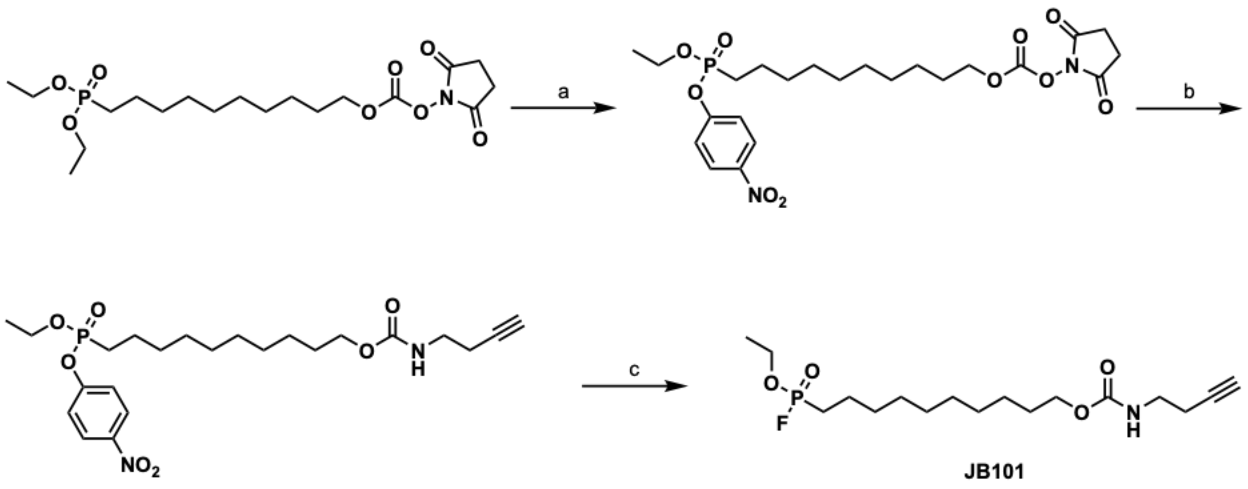


**Scheme S1**. Synthesis of JB101. Reagents and conditions: (a) 4-Nitrophenol, $(CF_3SO_2)_2O$, pyridine, $CH_2Cl_2$, 36%; (b) But-3-yn-1-amine, triethylamine, DMF, 49%; (c) Fluoride on polymer support, $CH_3CN$, 67%.

**Synthesis of JB101**

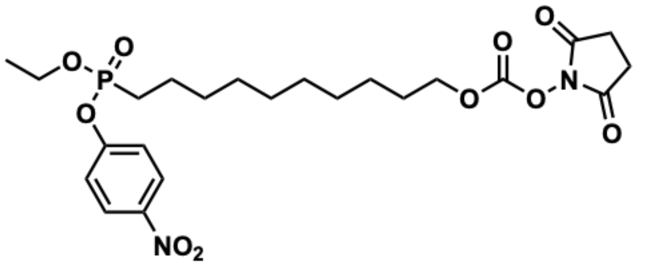

2,5-Dioxopyrrolidin-1-yl (10-(ethoxy(4-nitrophenoxy)phosphoryl)decyl) carbonate (**S1**)

To a stirred solution of 10-(diethoxyphosphoryl)decyl (2,5-dioxopyrrolidin-1-yl) carbonate(*46*) (100 mg, 0.230 mmol) in $CH_2Cl_2$ (1.25 mL) was added trifluoromethanesulfonic anhydride (53.0 µL, 0.315 mmol) immediately followed by pyridine (35.0 µL, 0.434 mmol). After stirring at ambient temperature for 10 minutes, 4-nitrophenol (48.0 mg, 0.345 mmol) was added to the reaction mixture. After stirring for an additional 30 minutes, the reaction mixture was concentrated in vacuo. The residue was purified by flash column chromatography on silica gel (gradient 35% → 100% ethyl acetate in hexane) to afford 44 mg (36%) of mixed phosphonate **S1** as a colorless oil. Obtained **S1** was directly used for the next step.

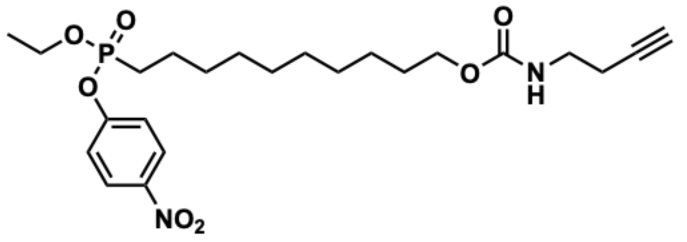

10-(Ethoxy(4-nitrophenoxy)phosphoryl)decyl but-3-yn-1-ylcarbamate (**S2**)

To a stirred solution of **S1** (30.0 mg, 94.4 µmol) and but-3-yn-1-amine (57.8 mg, 133 µmol) in DMF (2.0 mL) was added triethylamine (0.910 mg, 9.00 µmol). After being stirred for 20 minutes at ambient temperature, the solvent was removed under high vacuum. The residue was dissolved in $CH_2Cl_2$ and washed with saturated $NH_4Cl$, water, and brine. The organic layer was dried over $Na_2SO_4$ and concentrated in vacuo. The resulting residue was purified by flash column chromatography on silica gel (gradient 0% → 100% ethyl acetate in hexane) to afford 13.3 mg (49%) of **S2** as a colorless oil: $^1$H NMR (400 MHz, $CDCl_3$) δ 8.23 (d, *J* = 7.2 Hz, 2H), 7.38 (d, *J* = 9.1 Hz, 2H), 4.98 (s, 1H), 4.28 – 4.19 (m, 1H), 4.04 (t, *J* = 6.9 Hz, 2H), 3.36 – 3.30 (m, 2H), 2.42 – 2.37 (m, 2H), 1.97 – 1.88 (m, 2H), 1.73 – 1.55 (m, 4H), 1.47 – 1.34 (m, 3H), 1.33 – 1.24 (m, 14H); $^{31}$P NMR (162 MHz, $CDCl_3$): δ 30.48.

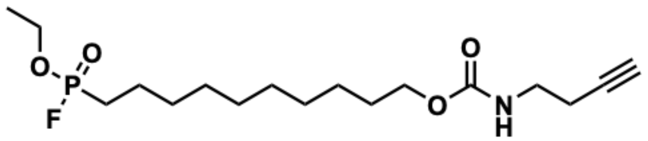

10-(Ethoxyfluorophosphoryl)decyl but-3-yn-1-ylcarbamate (**JB101**)

Fluoride on polymer support reagent (28.5 mg, 2.0 mmol/g) was added to a solution of **S2** (13.3 mg, 27.6 µmol) in $CH_3CN$ (250 µL). After being stirred for 2 hours, the supernatant was filtered through a disposable syringe with 0.2 µM filter. The resin was washed with $CH_2Cl_2/CH_3CN$ (2:1, v/v, 1.5 mL) by 5 minutes under stirring and the supernatant was collected by the same filtration. The combined filtrates were concentrated under reduced pressure, redissolved in $H_2O/CH_3CN$ (1:1, v/v, 1.0 mL), and lyophilized to afford 6.7 mg (67%) of **JB101** as a colorless oil: $^1$H NMR (400 MHz, DMSO-*d*$_6$) δ 7.20 (s, 1H), 4.17 (quint, *J* = 7.0 Hz, 2H), 3.91 (t, *J* = 6.7 Hz, 2H), 3.07 (q, *J* = 7.2 Hz, 2H), 2.80 (m, 1H), 2.29 – 2.25 (m, 2H), 2.08 – 1.92 (m, 2H), 1.57 – 1.46 (m, 4H), 1.36 – 1.24 (m, 15H); $^{19}$F NMR (376 MHz, DMSO-*d*$_6$): δ -62.64 (d, *J* = 1059.5 Hz); $^{31}$P NMR (162 MHz, DMSO-*d*$_6$): δ 32.61 (d, *J* = 1060.9 Hz).

• $^{1}$H NMR spectrum of compound S2 ($CDCl_3$, 400 MHz)

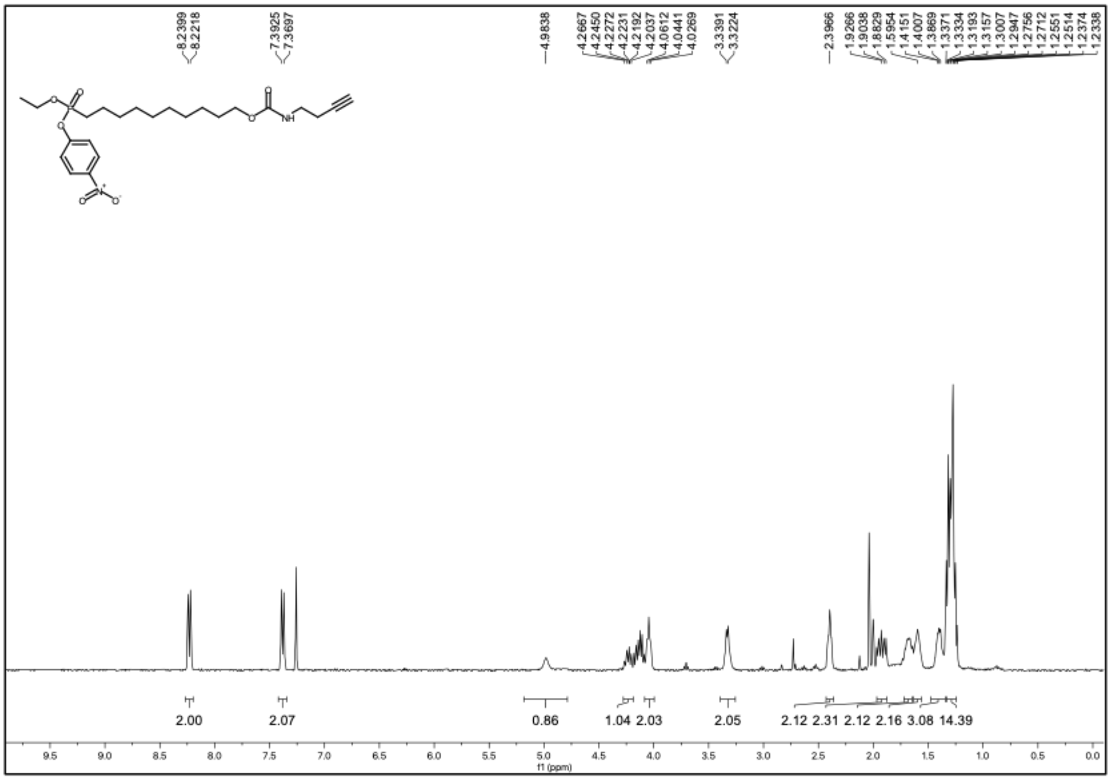

• $^{31}$P NMR spectrum of compound S2 ($CDCl_3$, 162 MHz)

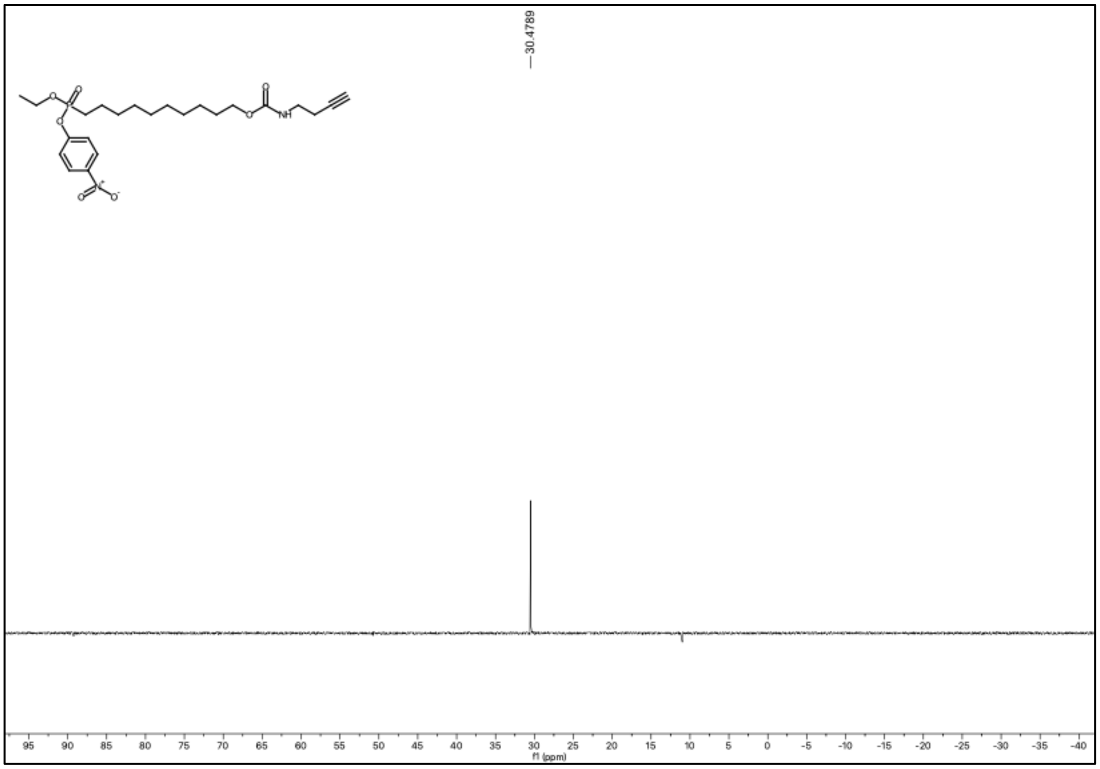

• $^{1}$H NMR spectrum of compound JB101 (DMSO-$d_6$, 400 MHz)

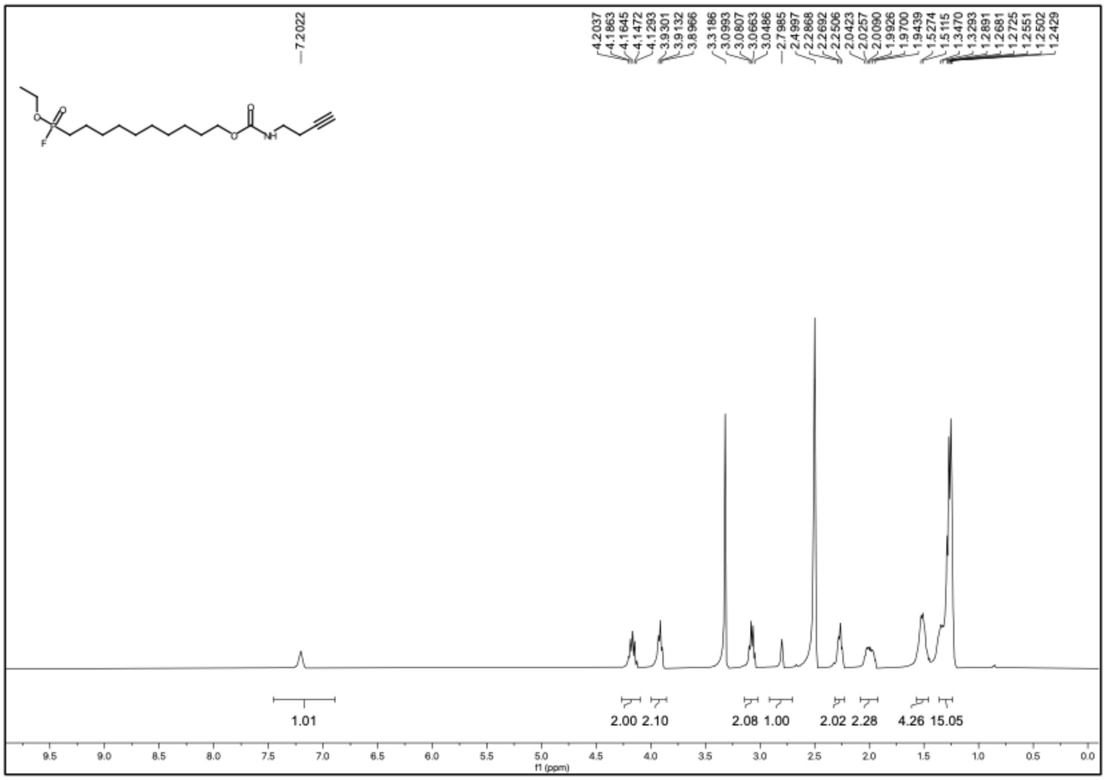


• $^{19}$F NMR spectrum of compound JB101 (DMSO-$d_6$, 376 MHz)

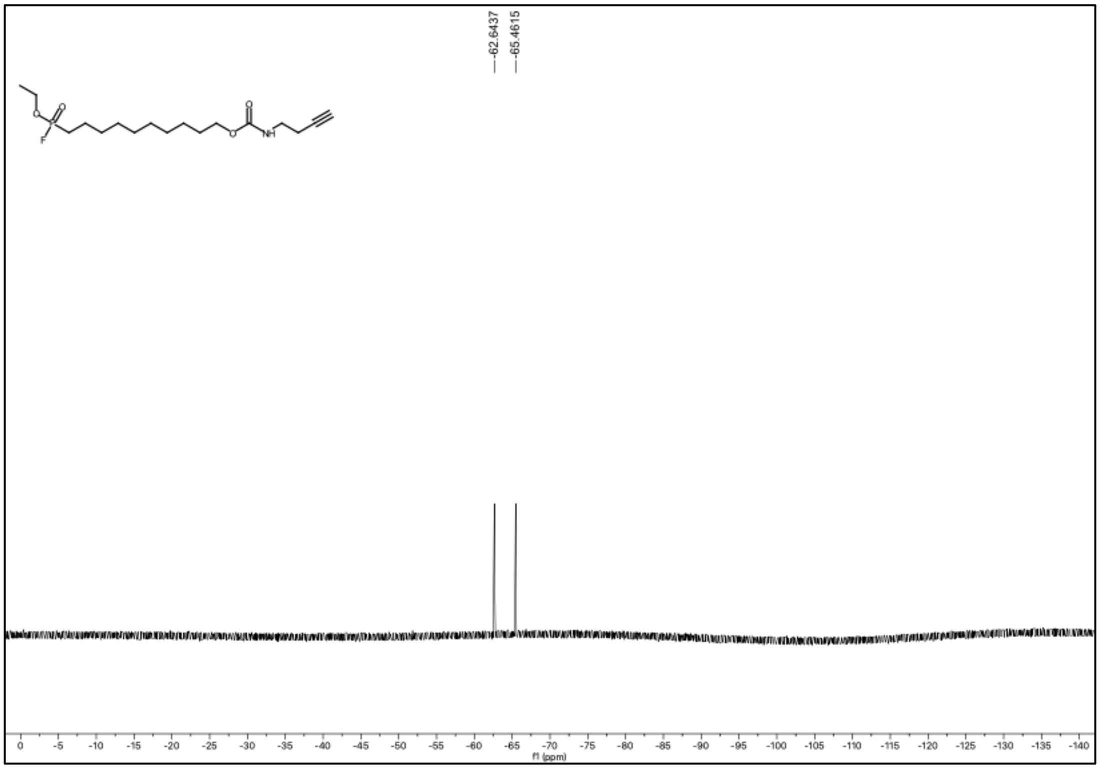

• $^{31}$P NMR spectrum of compound JB101 (DMSO-$d_6$, 162 MHz)

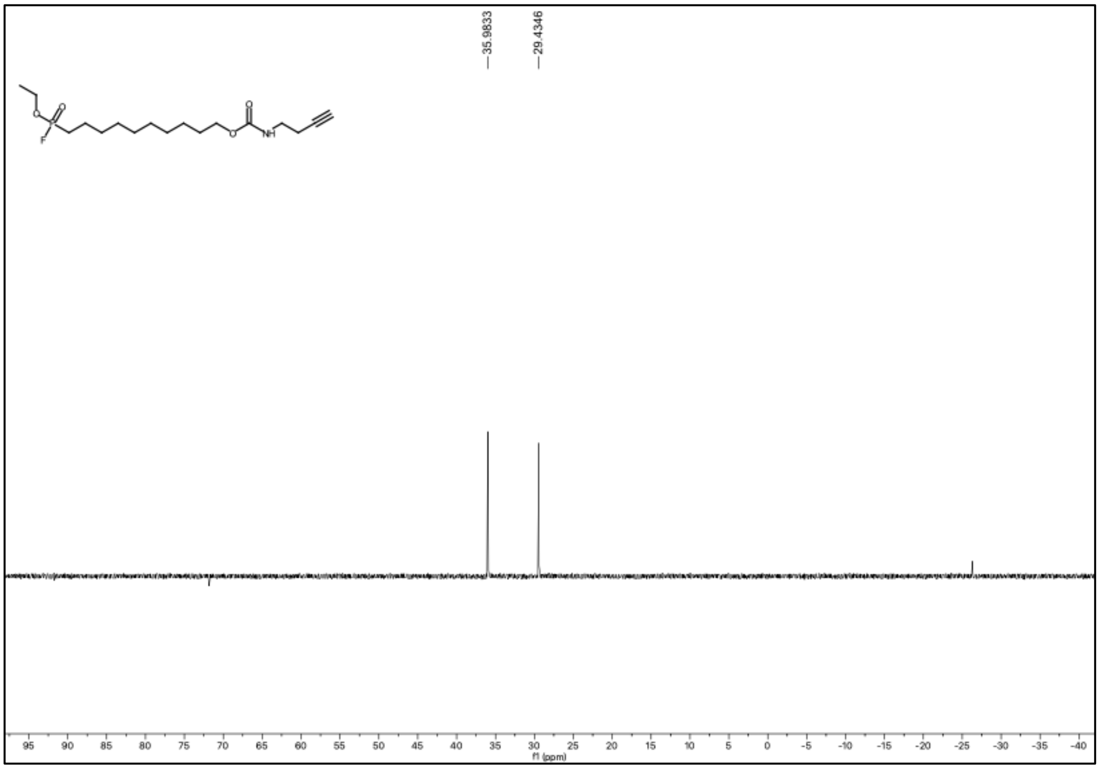

### References


1. M. Varadi *et al.*, AlphaFold Protein Structure Database: massively expanding the structural coverage of protein-sequence space with high-accuracy models. *Nucleic acids research* **50**, D439-D444 (2022).
2. Y. Fan *et al.*, The catalytic mechanism of direction-dependent interactions for 2, 3-dihydroxybenzoate decarboxylase. *Applied Microbiology and Biotechnology* **107**, 7451-7462 (2023).
3. X. Gao *et al.*, Structural basis of salicylic acid decarboxylase reveals a unique substrate recognition mode and access channel. *Journal of Agricultural and Food Chemistry* **69**, 11616-11625 (2021).
4. M. Song *et al.*, 2, 3-Dihydroxybenzoic Acid Decarboxylase from Fusarium oxysporum: Crystal Structures and Substrate Recognition Mechanism. *ChemBioChem* **21**, 2950-2956 (2020).
5. J. Jo *et al.*, Development of Oxadiazolone Activity-Based Probes Targeting FphE for Specific Detection of Staphylococcus aureus Infections. *J Am Chem Soc* **146**, 6880-6892 (2024).
6. T. Upadhyay *et al.*, Identification of covalent inhibitors of Staphylococcus aureus serine hydrolases important for virulence and biofilm formation. *Nature Communications* **16**, 5046 (2025).
7. A. T. Bakker *et al.*, Chemical Proteomics Reveals Antibiotic Targets of Oxadiazolones in MRSA. *J Am Chem Soc* **145**, 1136-1143 (2023).
8. M. H. Wei, University of Pennsylvania, (2023).
9. B. D. Saylor, J. J. Love, A secreted Staphylococcus aureus lipase engineered for enhanced alcohol affinity for fatty acid esterification. *Journal of Molecular Catalysis B: Enzymatic* **133**, S44-S52 (2016).
10. A. Ray, K. A. Edmonds, L. D. Palmer, E. P. Skaar, D. P. Giedroc, Glucose-Induced Biofilm Accessory Protein A (GbaA) Is a Monothiol-Dependent Electrophile Sensor. *Biochemistry* **59**, 2882-2895 (2020).
11. X. Xiao, Y. Li, L. Li, Y. Q. Xiong, Identification of Methicillin-Resistant Staphylococcus aureus (MRSA) Genetic Factors Involved in Human Endothelial Cells Damage, an Important Phenotype Correlated with Persistent Endovascular Infection. *Antibiotics* **11**, 316 (2022).
12. C. S. Lentz *et al.*, Identification of a *S. aureus* virulence factor by activity-based protein profiling (ABPP). *Nat Chem Biol* **14**, 609-617 (2018).
13. M. Fellner *et al.*, Structural Basis for the Inhibitor and Substrate Specificity of the Unique Fph Serine Hydrolases of Staphylococcus aureus. *ACS Infect Dis* **6**, 2771-2782 (2020).
14. L. Yu *et al.*, A Novel Repressor of the ica Locus Discovered in Clinically Isolated Super-Biofilm-Elaborating Staphylococcus aureus. *mBio* **8**, (2017).
15. C. J. Caballero *et al.*, The regulon of the RNA chaperone CspA and its auto-regulation in Staphylococcus aureus. *Nucleic Acids Res* **46**, 1345-1361 (2018).
16. S. Michalik *et al.*, A global Staphylococcus aureus proteome resource applied to the in vivo characterization of host-pathogen interactions. *Sci Rep* **7**, 9718 (2017).
17. M. Fellner, Newly discovered Staphylococcus aureus serine hydrolase probe and drug targets. *ADMET DMPK* **10**, 107-114 (2022).

18. R. Monteiro *et al.*, Proteome of a methicillin-resistant Staphylococcus aureus clinical strain of sequence type ST398. *J Proteomics* **75**, 2892-2915 (2012).
19. S. Reiss *et al.*, Global analysis of the Staphylococcus aureus response to mupirocin. *Antimicrob Agents Chemother* **56**, 787-804 (2012).
20. J. A. Cuaron *et al.*, Tea tree oil-induced transcriptional alterations in Staphylococcus aureus. *Phytother Res* **27**, 390-396 (2013).
21. S. Fuchs *et al.*, Aureolib - a proteome signature library: towards an understanding of staphylococcus aureus pathophysiology. *PLoS One* **8**, e70669 (2013).
22. T. N. B. Le, (2008).
23. C. Kohler *et al.*, A defect in menadione biosynthesis induces global changes in gene expression in Staphylococcus aureus. *J Bacteriol* **190**, 6351-6364 (2008).
24. J. T. Riordan *et al.*, Alterations in the transcriptome and antibiotic susceptibility of Staphylococcus aureus grown in the presence of diclofenac. *Ann Clin Microbiol Antimicrob* **10**, 30 (2011).
25. D. Becher *et al.*, A proteomic view of an important human pathogen--towards the quantification of the entire Staphylococcus aureus proteome. *PLoS One* **4**, e8176 (2009).
26. D. S. Ko *et al.*, Comparative genomics of bovine mastitis-origin Staphylococcus aureus strains classified into prevalent human genotypes. *Res Vet Sci* **139**, 67-77 (2021).
27. H. Peng *et al.*, Sulfide Homeostasis and Nitroxyl Intersect via Formation of Reactive Sulfur Species in Staphylococcus aureus. *mSphere* **2**, (2017).
28. C. Kohler *et al.*, Proteomic and Membrane Lipid Correlates of Re-duced Host Defense Peptide Susceptibility in a snoD Mutant of Staphylococcus aureus. *Antibiotics-Basel* **8**, (2019).
29. D. Frees *et al.*, New Insights into Stress Tolerance and Virulence Regulation from an Analysis of the Role of the ClpP Protease in the Strains Newman, COL, and SA564. *J Proteome Res* **11**, 95-108 (2012).
30. A. Jansen *et al.*, Production of capsular polysaccharide does not influence vancomycin susceptibility. *Bmc Microbiology* **13**, (2013).
31. M. Bischoff *et al.*, Microarray-based analysis of the σ regulon. *Journal of Bacteriology* **186**, 4085-4099 (2004).
32. K. E. Beenken *et al.*, Global gene expression in Staphylococcus aureus biofilms. *J Bacteriol* **186**, 4665-4684 (2004).
33. C. Burlak *et al.*, Global analysis of community-associated methicillin-resistant Staphylococcus aureus exoproteins reveals molecules produced in vitro and during infection. *Cell Microbiol* **9**, 1172-1190 (2007).
34. I. Staub, S. A. Sieber, beta-Lactam Probes As Selective Chemical-Proteomic Tools for the Identification and Functional Characterization of Resistance Associated Enzymes in MRSA. *Journal of the American Chemical Society* **131**, 6271-6276 (2009).
35. L. O. Henderson *et al.*, Transcriptional profiling of the L. monocytogenes PrfA regulon identifies six novel putative PrfA-regulated genes. *FEMS Microbiol Lett* **367**, (2020).
36. L. Chen, L. J. Keller, E. Cordasco, M. Bogyo, C. S. Lentz, Fluorescent triazole urea activity-based probes for the single-cell phenotypic characterization of *Staphylococcus aureus*. *Angew Chem Int Ed Engl* **58**, 5643-5647 (2019).
37. W. Sianglum, P. Srimanote, P. W. Taylor, H. Rosado, S. P. Voravuthikunchai, Transcriptome analysis of responses to rhodomyrtone in methicillin-resistant Staphylococcus aureus. *PLoS One* **7**, e45744 (2012).

38. A. K. Marr *et al.*, Overexpression of PrfA leads to growth inhibition of in glucose-containing culture media by interfering with glucose uptake. *Journal of Bacteriology* **188**, 3887-3901 (2006).
39. S. S. Chatterjee *et al.*, Intracellular gene expression profile of Listeria monocytogenes. *Infect Immun* **74**, 1323-1338 (2006).
40. M. Begley, R. D. Sleator, C. G. Gahan, C. Hill, Contribution of three bile-associated loci, bsh, pva, and btlB, to gastrointestinal persistence and bile tolerance of Listeria monocytogenes. *Infect Immun* **73**, 894-904 (2005).
41. C. Becavin *et al.*, Listeriomics: an Interactive Web Platform for Systems Biology of Listeria. *mSystems* **2**, (2017).
42. S. Wang *et al.*, An mRNA Display Approach for Covalent Targeting of a Staphylococcus aureus Virulence Factor. *Journal of the American Chemical Society* **147**, 8312-8325 (2025).
43. L. Holm, A. Laiho, P. Toronen, M. Salgado, DALI shines a light on remote homologs: One hundred discoveries. *Protein Sci* **32**, e4519 (2023).
44. C. UniProt, UniProt: the Universal Protein Knowledgebase in 2023. *Nucleic Acids Res* **51**, D523-D531 (2023).
45. J. Jumper *et al.*, Highly accurate protein structure prediction with AlphaFold. *Nature* **596**, 583-589 (2021).
46. S. E. Tully, B. F. Cravatt, Activity-Based Probes That Target Functional Subclasses of Phospholipases in Proteomes. *Journal of the American Chemical Society* **132**, 3264-3265 (2010).
